# The information bottleneck as a principle underlying multi-area cortical representations during decision-making

**DOI:** 10.1101/2023.07.12.548742

**Authors:** Michael Kleinman, Tian Wang, Derek Xiao, Ebrahim Feghhi, Kenji Lee, Nicole Carr, Yuke Li, Nima Hadidi, Chandramouli Chandrasekaran, Jonathan C. Kao

## Abstract

Decision-making emerges from distributed computations across multiple brain areas, but it is unclear *why* the brain distributes the computation. In deep learning, artificial neural networks use multiple areas (or layers) and form optimal representations of task inputs. These optimal representations are *sufficient* to perform the task well, but *minimal* so they are invariant to other irrelevant variables. We recorded single neurons and multiunits in dorsolateral prefrontal cortex (DLPFC) and dorsal premotor cortex (PMd) in monkeys during a perceptual decision-making task. We found that while DLPFC represents task-related inputs required to compute the choice, the downstream PMd contains a minimal sufficient, or optimal, representation of the choice. To identify a mechanism for how cortex may form these optimal representations, we trained a multi-area recurrent neural network (RNN) to perform the task. Remarkably, DLPFC and PMd resembling representations emerged in the early and late areas of the multi-area RNN, respectively. The DLPFC-resembling area partially orthogonalized choice information and task inputs and this choice information was preferentially propagated to downstream areas through selective alignment with inter-area connections, while remaining task information was not. Our results suggest that cortex uses multi-area computation to form minimal sufficient representations by preferential propagation of relevant information between areas.

**Significance:** The brain uses multiple areas for cognition, decision-making, and action, but it is unclear why cortical activity differs by brain area. Machine learning and information theory suggests that one benefit of multiple areas is that it provides an “information bottleneck” that compresses inputs into an optimal representation that is minimal and sufficient to solve the task. Combining experimental recordings from behaving animals and computational simulations, we show that later brain areas have a tendency to form such minimal sufficient representations of task inputs through preferential propagation of task-relevant information present in earlier areas. Our results thus provide insight into one possible reason why the brain uses multiple brain areas for supporting decision-making and action.

## Introduction

The brain uses multiple areas to perform cognitive functions and tasks, including decision-making, multisensory integration, attention, motor control, and timing ^1–10^. However, we lack a principled understanding of how and why computations and representations differ by brain area. For example, is all stimuli and decision-related information present in all brain areas^11,12^, or do the cortical representations differ depending on their processing stage ^13^? If the representations differ, are there general principles that can explain why the cortical representations differ by brain area?

To answer these questions, we draw on the *information bottleneck* (IB) *principle* from Machine Learning and Information Theory. The IB principle defines an *optimal* representation as a representation that is minimal and sufficient for a task or set of tasks ^14–16^. To understand this principle, consider the binary classification task in Fig. 1a, where the task *Y* is to answer if an image, **x**, is a dog. To answer this question correctly, we do not need to store every pixel of the image, but rather can form a compressed representation of the input. More generally, for a representation **z** to be useful (we define a representation **z** to be a function of the input **x**, that is **z** = *f* (**x**)), it should be sufficient for performing the task, containing similar *task* information to the input itself (mathematically, *I*(**Z**; *Y*) ≈ *I*(**X**; *Y*), where *I* denotes the mutual information). Among all such representations, the IB defines an optimal representation to be one that is also maximally compressed, or contains minimal information *I*(**Z**; **X**) about the input. Such minimal sufficient representations are proven to be robust to nuisance input variability unrelated to the task, such as the color of the dog or the background ^16^. Although these representations contain less information than the input, they are often more useful and robust representations for solving the task ^16–18^.

**Figure 1:**
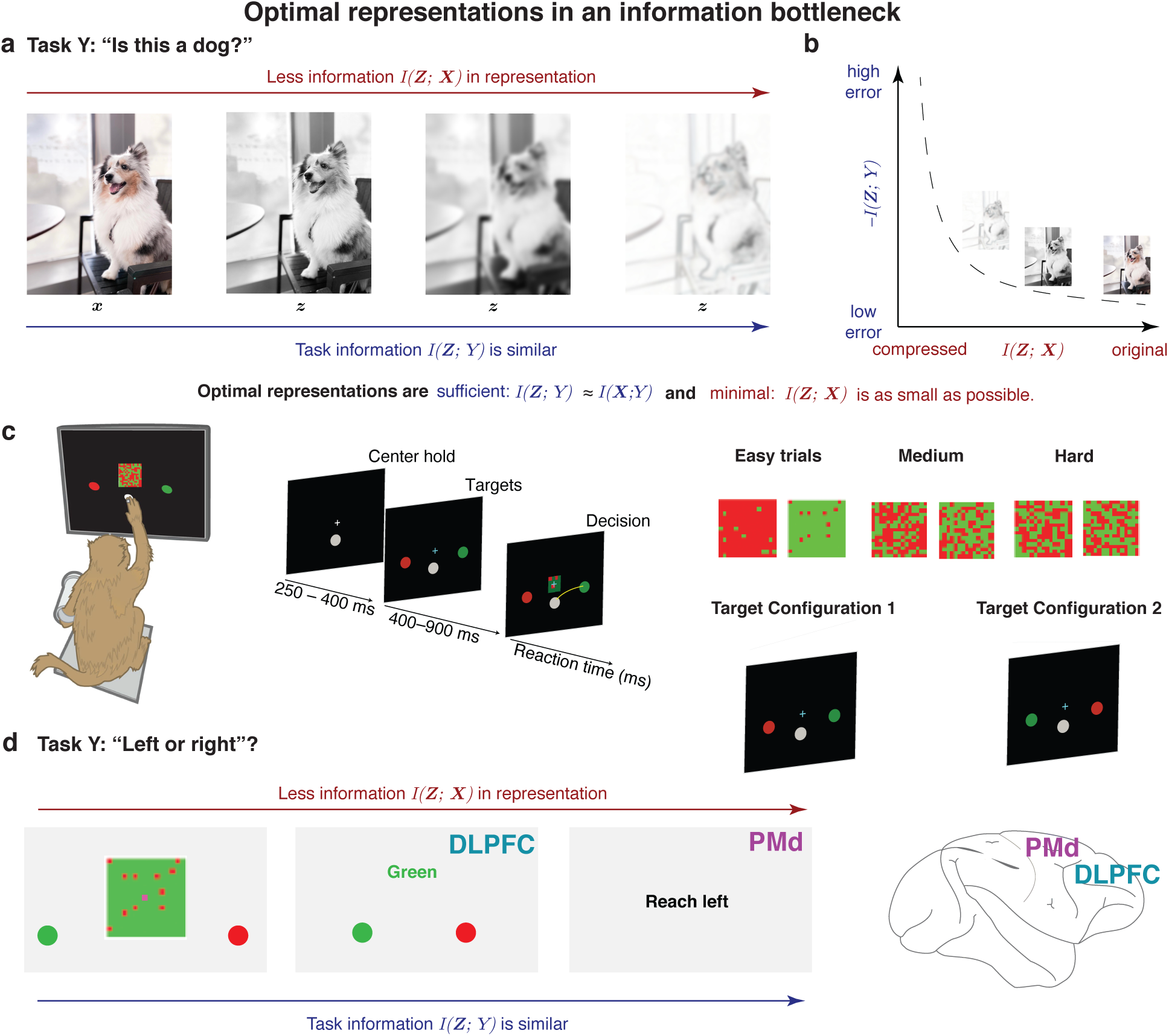
Optimal representations are formed through an information bottleneck. **(a)** Consider the task *Y* of discerning whether the image is of a dog. Images to the right have less information than the original image (*I*(**Z**; **X**) is smaller) but still contain approximately the same amount of information, *I*(**Z**; *Y*) to perform the task: “is this a dog?” **(b)** The information bottleneck trades off minimality, *I*(**Z**; **X**) as small as possible, with sufficiency, *I*(**Z**; *Y*) *≈ I*(**X**; *Y*). **(c)** Checkerboard task. The monkey reaches to the target whose color matches the checkerboard dominant color. Because there are two equally likely target configurations where the color of the left and right targets are swapped, this task unmixes the color and direction choice. **(d)** A minimal sufficient representation of this task is to only retain the direction decision, in this case, reach left. A cortical information bottleneck should therefore only find direction choice information in premotor and motor output areas.

An empirical observation from Machine Learning is that deep neural networks tend to form minimal sufficient representations in the last layers. Although multi-layer computation is not necessary for an IB, they provide a sufficient and even “natural” way to form an IB. A representation **z** = *f* (**x**) cannot contain more information than the input **x** itself due to the data processing inequality ^19^. Thus, adding additional layers typically results in representations that contain less information about the input. This is illustrated in Fig. 1a, where different transformations of the original image decrease the mutual information *I*(**Z**; **X**) in the representation. But the task information contained in the representation, *I*(**Z**; *Y*), is similar, since all images can be used to correctly answer “is this a dog?” (Fig. 1b) In visual cortex, which also has a hierarchical structure of computations, early layers of processing have representations that contain low-level details (e.g., edges) and deeper layers represent higher-level concepts (e.g., object identity) ^20,21^, suggesting that these areas subserve different functions or tasks.

But cortex may not necessarily implement an IB. The alternative hypothesis to IB is that the cortex does not form minimal sufficient representations. One manifestation of this alternative hypothesis is the “InfoMax” principle, where downstream representations are not minimal but rather contain maximal input information ^22^. This means information about task inputs not required to perform the task are present in downstream output areas. Two potential benefits of an InfoMax principle are (1) to increase redundancy in cortical areas and thereby provide fault tolerance, and (2) for each area to support a wide variety of tasks and thereby improve the ability of brain areas to guide many different behaviors.

In contrast to InfoMax, the IB principle makes two testable predictions about cortical representations. **Prediction 1:** there exists a downstream area of cortex that has a minimal and sufficient representation to perform a task (i.e., *I*(*X*; *Z*) is minimal while preserving task information so that *I*(*Z*; *Y*) ≈ *I*(*X*; *Y*)). **Prediction 2 (corollary if Prediction 1 is true):** there exists an upstream area of cortex that has more task information than the minimal sufficient area.

We tested these hypotheses by combining electrophysiological recordings in behaving monkeys from multiple brain areas, and modeling using recurrent neural networks during a perceptual decision-making task. In particular, we recorded from the dorsoateral prefrontal cortex (DLPFC) and dorsal premotor cortex (PMd) as monkeys performed a decision-making task called the Checkerboard Task (Fig. 1c). In this task, the monkey discriminated the dominant color of a checkerboard composed of red and green squares and reached to a target matching the dominant color. Because the red and green target locations were randomly assigned to be left or right on each trial (“target configuration”), the direction decision is independent of the color decision. That is, a green color decision is equally likely to correspond to a left or right decision. The animal’s behavioral report was either a right or left reach, determined after combining the sensory evidence with the target configuration (Fig. 1d). While color is initially needed to solve the task, the minimal sufficient representation of the task to generate the correct output is a representation of only the direction decision without the color decision or the target configuration. The IB principle would therefore predict that (1) downstream areas should only contain the direction choice (in Fig. 1d, “reach left”) and (2) upstream areas should contain information about the task inputs and decision-making process, including the target configuration, perceived dominant color of the checkerboard, and direction choice.

Consistent with these predictions of the IB principle, we found that DLPFC has information about the color, target configuration, and direction. In contrast, PMd had a minimal sufficient representation of the direction choice. Our recordings therefore identified a cortical IB. However, we emphasize the IB does not tell us *where* or *how* the minimal sufficient representation is formed. Instead, only our empirical results implicate DLPFC-PMd in an IB computation. Further, to propose a mechanism for how this IB is formed, we trained a multi-area RNN to perform this task. We found that the RNN reproduced key features of DLPFC and PMd activity, enabling us to propose a mechanism for how cortex uses multiple areas to compute a minimal sufficient representation.

## Results

### The checkerboard task involves multiple brain areas

We trained monkeys to discriminate the dominant color of a central static checkerboard (15 × 15 grid) composed of red and green squares (Fig. 1c). The number of red and green squares was random on each trial, leading to different levels of discrimination difficulty. The signed color coherence linearly indicates which color is dominant in the checkerboard, with −1 corresponding to completely green, 0 to equal numbers of red and green squares, and +1 to completely red. If the monkey reached to the target matching the dominant checkerboard color, the trial was counted as a success and the monkey received a juice reward. Critically, the color decision (red or green) was decoupled from the direction decision (left or right) because the left and right target identities were random on every trial. Fig. 2a,b shows the monkey’s psychometric and reaction time (RT) behavior on this task. Monkeys made more errors and reacted more slowly for more ambiguous checkerboards compared to the almost completely red or completely green checkerboards ^23^.

**Figure 2:**
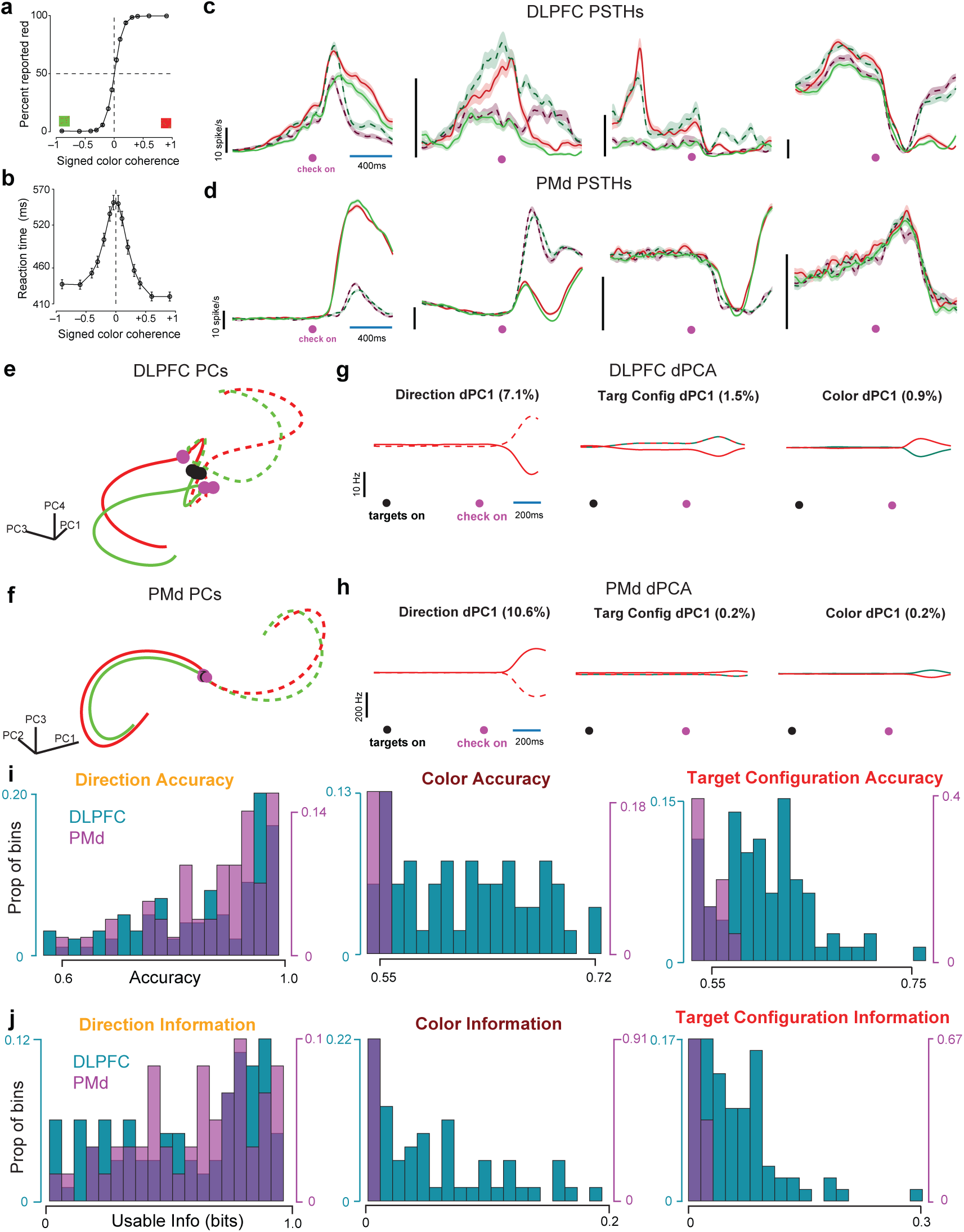
DLPFC and PMd recordings during the checkerboard task. **(a)** Psychometric and **(b)** reaction time curves of the monkey. X-axes in both (a) and (b) depict signed coherence which indicates the relative amount of red vs. green in the checkerboard. **(c)** Example DLPFC and **(d)** PMd PSTHs aligned to checkerboard onset. Red and green traces correspond to red and green color choices, respectively. Dotted and solid traces correspond to right and left direction choices, respectively. Data are smoothed with a 25 ms Gaussian and averaged across trials. **(e)** PCs 1, 3, and 4 for DLPFC and **(f)** PCs 1,2,3 for PMd. **(g)** Results of dPCA analysis for DLPFC and **(h)** PMd showing the dPCs for direction, target configuration, and color. **(i)** Histogram (across sessions) of direction, color, and target configuration decode accuracy and **(j)** usable information for DLPFC and PMd. The large variance in recordings is due to across-session variance.

We used linear multi-contact electrodes (U and V-probes) to record from the DLPFC (2819 single neurons and multiunits) and PMd (996 single neurons and multiunits) as the monkeys performed the checkerboard task. The PMd data was previously described in previous studies by Chandrasekaran and colleagues^23,24^. Example peri-stimulus time histograms (PSTHs) for neurons in DLPFC and PMd are shown in Fig. 2c, d and (and Fig. S5), respectively, where solid (dotted) lines correspond to left (right) reaches and color (red or green) denotes the color decision. DLPFC PSTHs in Fig. 2c separate based on direction choice, target configuration, and color choice, whereas PMd PSTHs primarily separate based on the direction choice, and only very modestly with target configuration or color.

Together, these examples demonstrate that DLPFC and PMd single units exhibit activity reflecting the decision-making process, implicating multiple brain areas in decision-making. Further, DLPFC likely contains multiple task-relevant signals, whereas PMd generally contains only direction choice related signals necessary for the behavioral report in the task. In the next sections, we use dimensionality reduction, decoding, and information theory to quantify the extent of color, target configuration, and direction representations in DLPFC and PMd at the population level and show that these physiological observations are consistent with the IB principle. We then use recurrent neural network models to build a mechanistic hypothesis for how an IB could be implemented.

### Evidence for a cortical information bottleneck between DLPFC and PMd

Our single neuron examples suggest that neuronal responses in DLPFC are modulated by color choice and target configuration, but PMd neurons generally are not. Our hypothesis is that these cortical representations in DLPFC and PMd are consistent with the IB principle. The direct prediction of this hypothesis is that the PMd population activity should contain a minimal and sufficient representation of the behaviorally relevant output – the direction choice – while upstream DLPFC population activity should represent multiple task-relevant variables.

To study this at the population level, we performed principal components analysis (PCA) on DLPFC and PMd neural population activity. In these PCA trajectories, we subtracted the condition-independent component of the signal to better highlight representations of direction, target configuration, and color. DLPFC and PMd exhibited qualitatively different neural population trajectories (Fig. 2e,f). First, after target onset, DLPFC trajectories separated as a function of the two target configurations, and at the time of checkerboard onset (purple dots), DLPFC activity further separated into four distinct trajectories based on the four possible color × direction outcomes (green left, green right, red left, red right, Fig. 2e). Thus, DLPFC contains information about target configuration, color choice, and direction choice. Note, we chose PC 1, 2 and 4 of DLPFC as they provided a better visualization of target configuration signal between target and checkerboard. In contrast, PMd trajectories in Fig. 2f did not exhibit target-configuration-specific steady state responses. Thus, at the point of checkerboard onset (purple dots), trajectories overlapped in the top 3 principal components, and only separated based on the direction, but not color.

To quantify these these differences, we performed demixed principal component analysis (dPCA) on the DLPFC and PMd population activity (Fig. 2g,h). DLPFC and PMd activity both exhibited strong condition independent activity (82% and 86% variance, respectively). DLPFC activity represented the target configuration, but PMd did not (Fig. 2g,h, target configuration dPC). dPCA also identified principal axes that maximized variance related to the direction choice, color choice, and target configuration. In DLPFC, the top direction choice, color choice, and target configuration axes captured 7.1%, 0.9%, and 1.5% of the population activity. In PMd, these values were 10.6%, 0.2%, and 0.2%. Across all direction, color, and target configuration axes, the dPCA variance captured for DLPFC was 11%, 3%, 5%, while for PMd it was 12%, 1%, 1%. This dPCA analysis provides further evidence that DLPFC represents direction, target configuration, and color while PMd has nearly minimal representations of color and target configuration, consistent with the IB principle. The difference in color and target configuration variance explained between DLPFC and PMd is statistically significant (*p <* 0.02, shuffle test shown in Fig. S6).

Our dPCA results with trial-averaged firing rates suggest that axes associated with color and target configuration in PMd have very little variance associated with them. However, these results do not rule out the possibility that there is decodable information about these task-related variables on single trials. We performed two other analyses to assess if there is an IB and that PMd contains a minimal sufficient representation. We calculated the decode accuracy and an estimate of mutual information for direction, color, and target configuration in DLPFC and PMd population activity. We decoded direction, color, and target configuration from DLPFC and PMd sessions where we recorded a small population of neurons using a support vector machine (see Methods). To estimate mutual information, we quantified the Usable Information, a variational lower bound to mutual information that can be computed on high-dimensional data through estimating cross-entropy loss ^17,18^ (see Methods).

We found that for many DLPFC sessions we could reliably decode direction, color, and target configuration on individual trials well above chance (histogram in Fig. 2i, mean decoding accuracy across sessions: direction 86%, target configuration 59%, color 57%, for details on decoding, see Methods). To conservatively assess significance, we shuffled the trials for each session 100 times and obtained a surrogate decoding accuracy for each session, and then tested if the true accuracy was greater than the 99^th^ percentile of the shuffled accuracy distribution. We found that 100%, 78%, 54% sessions having true accuracy higher than 99^th^ percentile of the shuffled accuracy for direction, target configuration and color. In DLPFC, we could therefore significantly decode direction choice, target configuration, and color choice. In contrast in PMd, we rarely found sessions where target configuration and color could be reliably decoded well above chance (mean accuracy across sessions: direction 88%, target configuration 52%, color 51%), as shown in Fig. 2i. Again, we compared the true decoding accuracy with the surrogate decoding accuracy obtained by shuffling trials. We found 98%, 26%, 19% sessions having true accuracy above 99^th^ percentile of the shuffled accuracy for direction, target configuration, and color. The differences in mean decoding accuracy in DLPFC and PMd were significant for only color and target configuration (*p <* 0.001, Wilcoxon rank-sum test), but not direction (*p* = 0.145, Wilcoxon rank-sum test). Thus, both DLPFC and PMd contain significantly more direction information but DLPFC contains more color choice and target configuration information consistent with the IB principle.

We also quantified usable information for DLPFC and PMd (Fig. 2j). For this analysis, we restricted it to sessions with significant decode accuracy with a session considered to have a significant decodability for a variable if the true accuracy was above the 99^th^ percentile of the shuffled accuracy for a session. This is because these sessions contain information about task variables. However, we also present the same analyses using all sessions in Fig. S7. DLPFC had sessions with non-zero usable information for direction, color, and target configuration (average direction information: 0.56 bits, target configuration: 0.07 bits, color: 0.06 bits) while PMd only had non-zero usable information for direction (average direction information: 0.64 bits, target configuration: 0.013 bits, color: 0.005-bits). The differences in usable information in DLPFC and PMd were also significant for only color and target configuration (*p <* 0.001, Wilcoxon rank-sum test), but not direction (*p* = 0.107, Wilcoxon rank-sum test). Together, these results indicate that PMd had a more minimal representation of task inputs, particularly color and target configuration, than DLPFC.

Our dPCA and decoding results are consistent with the existence of a cortical IB between DLPFC and PMd that reduces the amount of target configuration and color information in PMd while preserving the direction choice information necessary to solve the task. We next modeled this multi-area IB to develop a mechanistic hypothesis for how this cortical IB could be computationally implemented.

### A multi-area recurrent neural network model of DLPFC and PMd

To develop a mechanistic hypothesis for this cortical IB, we studied our previously reported multi-area RNN to perform the Checkerboard task (Fig. 3a)^25^. We chose to use this multi-area RNN because prior work demonstrated this RNN, like our PMd data, has a minimal color representation in Area 3. The RNN input was 4D representing the target configuration and checkerboard signed coherence, while the RNN output was 2D, representing decision variables for a left and right reach (see Methods). The RNN had 3 areas, obeyed Dale’s law ^26^, and had approximately 10% feedforward and 5% feedback connections between areas based on projections between prefrontal and premotor cortex in a macaque atlas^27^. RNN psychometric and RT curves for the multi-area RNN exhibited similar behavior to monkeys performing this task (Fig. 3b,c; across several RNNs, see Fig. S1).

**Figure 3:**
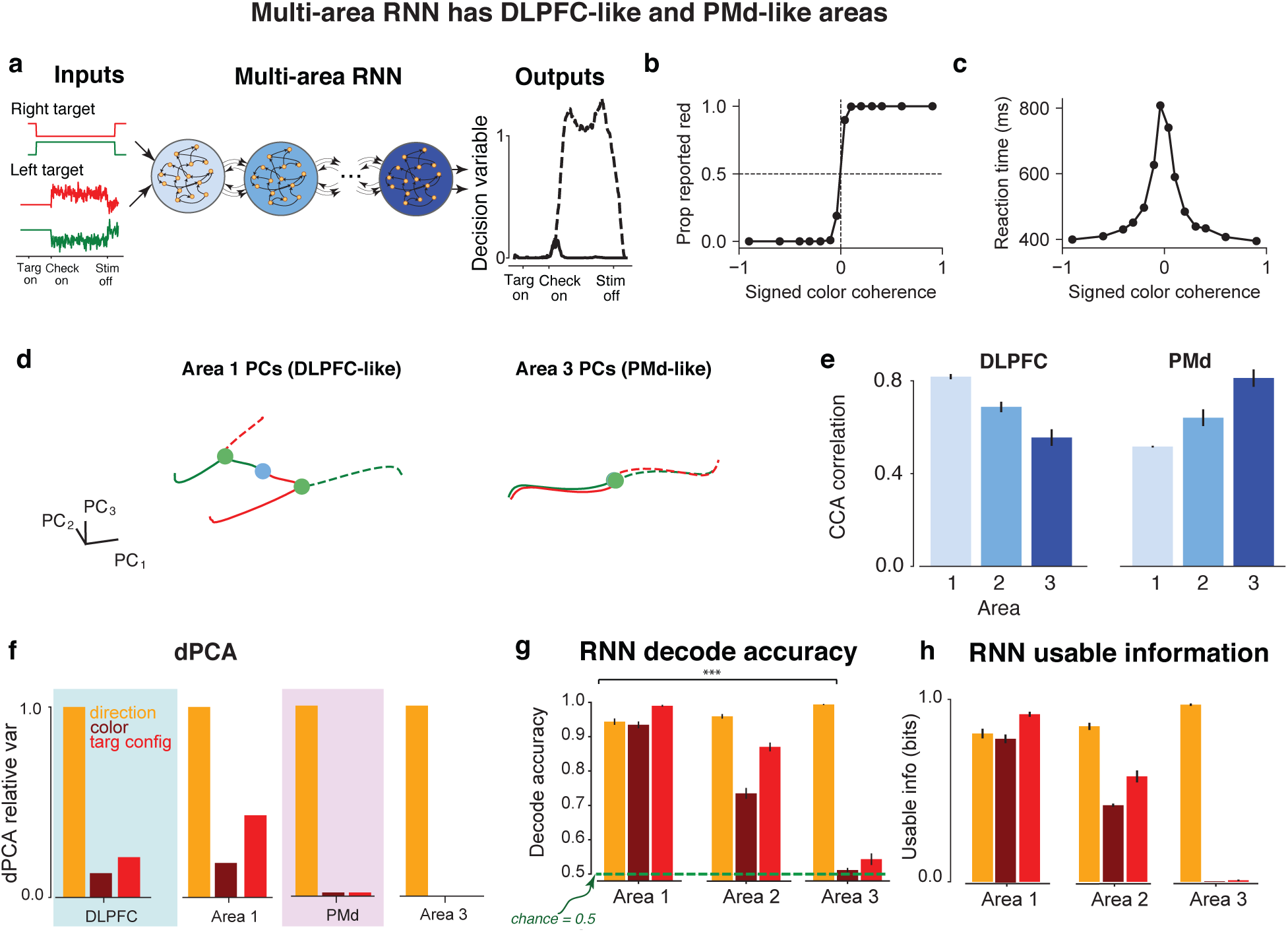
RNN modeling of the CB task. **(a)** Multi-area RNN configuration. The RNN received 4 inputs. The first two inputs indicated the identity of the left and right targets, which was red or green. These inputs were noiseless. The last two inputs indicated the value of the signed color coherence (proportional to amount of red in checkerboard) and negative signed color coherence (proportional to amount of green in checkerboard). We added independent Gaussian noise to these signals (see Methods). The network outputted two analog decision variables indicating evidence towards the right target (solid line) or left target (dashed line). A decision was made in the direction of whichever decision variable passed a preset threshold (0.6) first. The time at which the decision variable passed the threshold was defined to be the reaction time. **(b,c)** Psychometric and reaction time curves for exemplar multi-area RNN. **(d)** Area 1 and Area 3 principal components for exemplar RNN. **(e)** CCA correlation between each area and DLPFC principal components (left) and PMd principal components (right). DLPFC activity most strongly resembles Area 1, while PMd activity most strongly resembles Area 3. See also Fig. S3 where we computed CCA as a function of the number of dimensions. **(f)** Relative dPCA variance captured by the direction, color, and target configuration axes. Normalization makes direction variance equal to 1. Area 1 (3) variances more closely resemble DLPFC (PMd). **(g)** Area 1 has significantly higher decoding accuracies and **(h)** usable information compared to Area 3, consistent with DLPFC and PMd.

Remarkably, even though the multi-area RNN was in no way regularized to reproduce DLPFC and PMd activity, activity in Area 1 qualitatively resembled neural responses in DLPFC, representing both direction and color, while Area 3 resembled PMd, representing direction (Fig. 3d,e). Like DLPFC, Area 1 had four distinct trajectories corresponding to the four possible task outcomes and represented target configuration, direction choice, and color choice (Fig. 3d and see Fig. S2). In contrast, Area 3 population trajectories primarily separated based on direction and not by target configuration or color — remarkably similar to the trajectories observed in PMd.

We performed CCA to assess the similarity between the empirical neural trajectories to each RNN area’s neural trajectories (see Methods). The CCA analysis suggested that Area 1 exhibited the strongest resemblance to DLPFC, while Area 3 most strongly resembled PMd activity (Fig. 3e). These results show that a multi-area RNN reproduced similar behavior to the monkey, and further that it did so with architecturally and qualitatively distinct areas that resembled the physically distinct DLPFC and PMd cortical areas.

The RNN activity differs from cortical activity in two important ways. First, RNNs generally had a significantly smaller variance condition-independent signal (46.7% and 49.4% average variance in Area 1, and 3, respectively) than in DLPFC and PMd (82% and 86% variance, respectively). One possible explanation is that condition-independent variance in PMd is associated with a trigger signal, likely from the thalamus^28^, and these RNNs do not output arm kinematics, forces, or electromyography (EMG). Similarly, in DLPFC, we did not explicitly model the target and checkerboard inputs to have large onset signals that are often associated with visual stimulation. This significant condition independent variance in the neurophysiological data may therefore make decoding more difficult since there is relatively lower variance representing direction, color, or target configuration. While our CCA analysis was performed with the condition-independent signal removed, this difference impacts both dPCA and decoding results. In general, we found that RNN exhibited trends observed in the neurophysiological data more strongly, including more variance captured for direction, color, and target configuration, as well as higher decoding accuracies. We therefore compared the relative, rather than absolute, trends in RNN activity and DLPFC for dPCA and decoding analyses for the purposes of identifying a RNN IB similar to the cortical IB.

We found that the 3-area RNN exhibited similar trends to DLPFC and PMd activity in dPCA variance and decoding accuracy. When comparing only the top axis for direction, color, and target configuration, DLPFC activity had relatively large variance captured along the direction axis (7.1% variance captured), followed by relatively weaker representations for target configuration (1.5%) and color (0.9%). Area 1 activity had similar relative trends, with the direction axis explaining 30.9% variance, followed by target configuration (13.3%) and color (5.6%). In Fig. 3f, we show the similarity in relative trends in DLPFC and Area 1 by normalizing these quantities by the variance captured by the direction axis. Meanwhile, PMd activity exhibited more direction-related variance than DLPFC (10.6% variance captured) and did not exhibit a significant target configuration and color axis representation (0.2% for both axes). Likewise, Area 3 had a stronger representation of direction (48.5% variance captured) than in Area 1, but negligible target configuration and color axes variance (0.1% for both axes), demonstrating the same relative trend (Fig. 3f). These results show that Area 1 more strongly resembles DLPFC and Area 3 more strongly resembles PMd in relative dPCA variance.

We next evaluated the decode accuracy and usable information in the multi-area RNN using a nonlinear decoder on single trials, in line with prior work^29^ (see Methods). We found Area 1, like DLPFC, had significant information for direction, target configuration, and color. Direction, color, and target configuration could be decoded from Area 1 population activity at accuracies of 94.4%, 93.4%, and 99.0%, respectively, corresponding to 0.81, 0.79, and 0.92 bits of usable information (Fig. 3g,h). In contrast, Area 3 direction decode accuracy was 99.4%, while color and target configuration accuracy were significantly lower (51.1% and 54.3%, respectively). This corresponded to 0.97, 0.0023, and 0.0078 bits of usable information for direction, target configuration, and color, respectively. We also found analogous conclusions when using an SVM decoder (Fig. S4).

Together, these results show that our multi-area RNN exhibited distinct areas that resembled DLPFC and PMd activity, and also implemented an IB so that its output area only had primarily direction information and less color and target configuration information. This multi-area RNN therefore implements a candidate mechanism for how a cortical IB could be implemented between DLPFC and PMd.

### Mechanistic features of the DLPFC and PMd bottleneck: partial orthogonalization and selective propagation

The multi-area RNN contains representations consistent with the IB principle — its input and output areas resemble DLPFC and PMd. The RNN therefore models key aspects of our physiological data and is therefore a candidate system to understand how multiple areas could lead to the empirically observed minimal sufficient representations. The unique advantage of the RNN is that we know the firing rates in each area as well as the within and inter-areal connections, which allows us to investigate how the multi-area RNN deemphasizes color information through an IB. We reasoned that such an IB could be implemented in three ways: Color information may be (1) primarily attenuated through *recurrent neural dynamics*, (2) primarily attenuated through *inter-areal connections*, or (3) attenuated through a combination of recurrent dynamics and inter-areal connections. This is illustrated in Fig. 4a.

**Figure 4:**
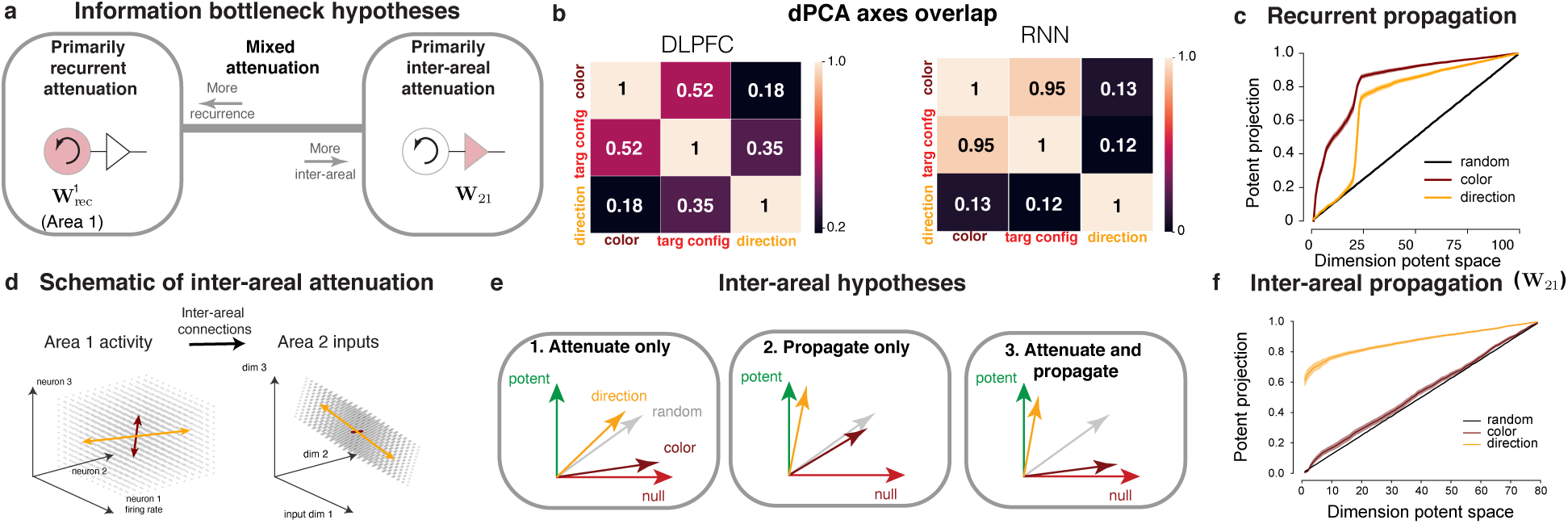
IB hypotheses and mechanism. **(a)** Candidate mechanisms for IB. **(b)** Axes overlap of the direction, color, and target configuration axes for DLPFC and RNN data. The direction axis is more orthogonal to the color and target configuration axes. **(c)** Projections onto the potent space of the intra-areal dynamics for Area 1 (for other areas, see Supp. Fig. S11). We computed the potent projection of the direction axis, color axis, and a random vector with each area’s intra-areal dynamics matrix. We found intra-areal dynamics amplify color information in Area 1, and do not selectively attenuate color information in Areas 2 and 3. **(d)** Illustration depicting how the orientation of the axes affect information propagation. Information on the direction axis (orange) can be selectively propagated through inter-areal connections which information on the color axis (maroon) is not. **(e)** Inter-areal hypotheses. **(f)** Projections onto the potent space between areas for the color axis, direction axis, and random vector. Regardless of the dimension of the potent space, the direction axis is preferentially aligned with the potent space, indicating the information along this axis propagates, while the color axis is approximately randomly aligned. We emphasize the high alignment of the direction axis: the direction axis has a stronger alignment onto the first potent dimension of **W**_21_ than the remaining dimensions combined. Meanwhile, the color axis is aligned at nearly chance levels, and will therefore be propagated significantly less than the direction axis. Shading indicates s.e.m.

To test these hypotheses, we first quantified how color, target configuration, and direction information was represented in the animals and our network. We performed dPCA in the different areas to identify demixed principal components that represented the corresponding information. In DLPFC, we quantified the overlap of the dPCA principal axes for target configuration, color, and direction. While the target configuration and color axes were relatively aligned (dot product, DP: 0.52), the direction axis was closer to orthogonal to the color axis (DP: 0.18) and the target configuration axis (DP: 0.35), as shown in Fig. 4b. These results suggest that DLPFC partially orthogonalizes information about the direction choice from the color choice and target configuration. We also observed these trends in the RNN, albeit more strongly. In DLPFC-resembling Area 1, we observed the target configuration and color were also highly aligned (DP: 0.95) but that the direction axis was more orthogonal to the color axis (DP: 0.13) and the target configuration axis (DP: 0.12). In our simulations, the reported values reflect the mean across 8 networks trained with the same hyperparameters. A candidate mechanism for this orthogonalization, found by performing dynamical analyses on the RNN, is shown in Fig. S8.

The advantage of our model is that both the intra-areal dynamics and inter-areal connectivity matrices are known. We analyzed how these axes were aligned with the intra-areal recurrent dynamics and inter-areal connectivity matrices to identify which hypothesis explained how the RNN implemented the IB. To do so, we performed singular value decomposition (SVD) on these matrices. We defined a *k*-dimensional “potent space” to be the right singular vectors corresponding to the *k* largest singular values of the matrix. The “null space” is the orthogonal complement of the potent space, which comprises the remaining *d* − *k* smallest singular vectors, where *d* refers to number of columns of the matrix. The null projection magnitudes are equal to one minus the potent projection. We quantified how the color and direction axis were aligned with these potent and null spaces (see Methods). This enabled us to study if the emergence of minimal sufficient representations was due to: (1) relative amplification of the direction information with respect to a random vector, (2) relative suppression of the color/target configuration information with respect to a random vector, or (3) a combination of both. Finally, we focused our analyses on Area 1 recurrent dynamics and the inter-areal connections between Areas 1 and 2 (**W**_21_) because color information is significantly attenuated by Area 2 (dPCA color variance in Area 2: 0.14%). The same analyses applied to downstream areas are shown in Fig. S11.

We first tested the hypothesis that the RNN IB is implemented primarily by recurrent dynamics (left side of Fig. 4a). These recurrent dynamics can be equivalently interpreted as the RNN implementing a feedforward neural network in time. We quantified how the color and direction axis were aligned with these potent and null spaces of the intra-areal recurrent dynamics matrix of Area 1 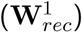. We did not include the target configuration axis for simplicity, since it highly aligns with the color axis for this network. The axes found through dPCA are proxies for how task information is represented in the neural activity over the trial, while the dynamical activity during a trial depends on the complex interaction between left and right singular vectors of the recurrent matrix (and task inputs) ^30^. We use the alignment of the (fixed) dPCA axes with the right singular vector as a proxy for the amount of information propagated through recurrence. In Area 1, we found significant alignment of the color axis with the top right singular vectors (potent space) of the recurrent dynamics matrix (Fig. 4c). Additionally, we also performed an alternative analysis where we compared input and activity representations of color discriminability and direction discriminability for our exemplar network. We observe an amplification, not a reduction, in color discriminability with respect to the inputs in Area 1 (Fig. S9) consistent with the amplification observed in Fig. 4c. These findings argue against the hypothesis that recurrent dynamics preferentially attenuate color information by projecting it into a nullspace of the recurrent dynamics. Rather, these data suggest that Area 1 has significant color information in its potent space, indicating that the recurrent computation amplifies color information. In Areas 2 and 3, the color axis (which had small variance of 0.14% and 0.07% in Areas 2 and 3, respectively) was again typically more strongly aligned with 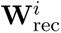 than a random vector, (Fig. S11). In summary, the dPCA and discriminability analyses suggest that the network did not use recurrent dynamics to attenuate color information, and is therefore inconsistent with the hypothesis that the IB is primarily implemented through intra-areal recurrent dynamics.

Our alternative hypothesis is that color information is primarily attenuated through inter-areal connections. This is schematized in Fig. 4d, where inter-areal connections propagate activity along the Area 1 direction axis (orange) to Area 2, but attenuate Area 1 color axis activity (maroon). To test this hypothesis, we quantified how the color and direction axis were aligned with these potent and null spaces of the inter-areal matrices. This enabled us to quantify the alignment of the direction and color axes with the inter-areal potent and null spaces and specifically determine how direction and color information were differentially propagated (Fig. 4e). Inter-areal connections could attenuate color information by aligning the color axis with the null space of **W**_21_ (Hypothesis 1 in Fig. 4e), propagate information preferentially (Hypothesis 2 in Fig. 4e), or both attenuate and propagate information (Hypothesis 3 in Fig. 4e).

We calculated the projections for both the color and choice axes on to the potent space for the connection matrix from area 1 to area 2 (**W**_21_) The projections onto the potent space are shown in Fig. 4f for the color and direction axis. We found the direction axis was more aligned with the potent space. In fact, the direction axis was consistently most aligned with the top singular vector of the **W**_21_ matrix, on average more than the remaining *d* − 1 singular vectors. In contrast, the color axis was aligned to a random vector. This suggests that learning in the multi-area recurrent network involved aligning the relevant information (in the activations) with the top singular vector (governed by the learned parameters of the feedforward matrix). These results indicate that direction information is preferentially propagated to subsequent areas, while color information is aligned with a random vector. This is most consistent with the “propagate only” hypothesis shown in Fig. 4e.

Such alignment of the direction axis with the top singular vector of the connection matrix isn’t trivial: the potent space depends on the parameter **W**_21_ learned during training, while the direction axis is not a parameter but a dPCA axis computed from Area 1 activity. This alignment was robust to the dimension of the effective potent space, and was consistent across networks with varying feedforward connectivity percentages (10%, 20%, 30%, 50%, 100%). Further, we found that **W**_21_ in unconstrained 3 area networks (i.e., without Dale’s law constraints) had significantly reduced alignment of the direction axis with the top singular vectors (Fig. S11d).

In summary, the multi-area RNN IB is primarily implemented through preferential propagation of direction information through inter-areal connections. Recurrent dynamics play a role in processing color information and target configuration to arrive at direction choice information. In Area 1, RNN dynamics amplify color information. Our results are therefore most consistent with the hypothesis that the IB is implemented primarily through inter-areal connections, not recurrent dynamics, in Fig. 4a.

### Effect of network architecture and training hyperparameters on the information bottleneck

We next assessed the network architectures and hyperparameters that influenced the formation of minimal sufficient representations during the Checkerboard task. We swept RNN architectural parameters and machine learning hyperparameters to assess what variables were important for learning minimal sufficient representations without color information. Specifically, we varied the connectivity type (unconstrained connectivity vs Dale’s law and varying proportion of connections), the percentage of unconstrained feedforward connections, the percentage of feedforward E to I connections, the percentage of E to E connections, the number of areas (from 1 to 4), the number of artificial networks, L2 weight regularization, L2 rate regularization, and the learning rate. Our choice of feedforward and feedback connections were informed by a macaque atlas^27^. In our sweeps we quantified the color (and direction) variance and accuracy in the last area.

We generally observed minimal sufficient representations in the last area so long as there was a sufficient *connection* bottleneck between RNN areas. In unconstrained networks, shown in Fig. 5a, color variance and decode accuracy decreased as the percentage of feedforward connections between areas decreased, though the representations were not minimal. We incorporated Dale’s law with 80% E and 20% I neurons following Song et al. ^26^ into subsequent sweeps (Fig. 5b-e). Minimal representations with chance color decode accuracy emerged when the percentage of feedforward E to I connections was 2 − 5% or less (the overall percentage of feedforward E to E was fixed at 10% following a macaque atlas). We also found that when there was no feedforward inhibition, but when we varied the percentage of feedforward or feedback E-to-E connections RNNs generally had nearly minimal representations (Fig. 5c, and Supp. Fig. S12). We observed that as long as there were 3 or 4 areas, there was a large decrease in color information in the last area (Fig. 5d) quantified by decoding, though note that there was a large drop in color variance for 2 area networks. These results suggest that multi-area networks, with a feedforward connection bottleneck tend to produce more minimal representations for the Checkerboard task.

**Figure 5:**
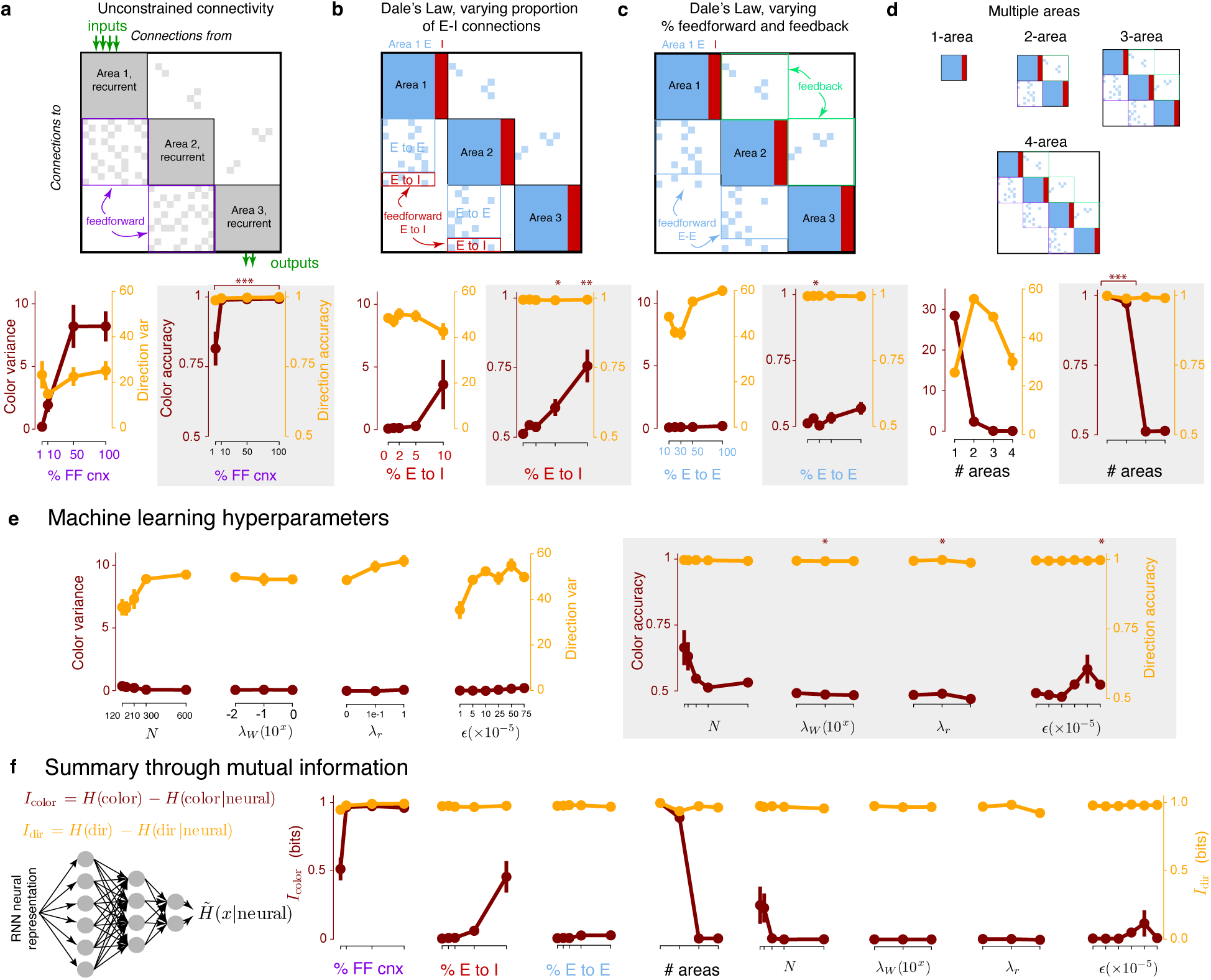
Robustness of the information bottleneck across hyperparameters and computational advantage. Varying **(a)** proportion of feedforward connections in an unconstrained network, **(b)** E-I connections in a Dale’s law network, **(c)** the proportion of feedforward E-E connections in a Dale’s law network; feedback connections are varied in Fig. S12, **(d)** the number of areas, and **(e)** the machine learning hyperparameters revealed that Area 3 color variance and color accuracy decrease as long as there is a connectivity bottleneck between areas. **(f)** Summary of these results quantifying usable information. Overall, neurophysiological architecture constraints in the form of multiple areas, sparser connections between areas than within an area, as well as a smaller fraction of E-I connections lead to a minimal color representation in the last area. Parameters of the exemplar network are in Table 1.

We also varied machine learning hyperparameters (Fig. 5e) to assess the extent to which the IB was present. To prevent an exponential search space, we fixed the architecture to the exemplar network used in this study and tested one hyperparameter at a time. We varied the number of artificial units in the network, the L2 weight regularization, the L2 rate regularization, and the learning rate. At each hyperparameter setting, we trained a total of 8 multi-area RNNs. Our exemplar network consistently exhibited little to no Area 3 color information across every hyperparameter setting we chose, suggesting that the presence of the IB is not a result of a particular choice of machine learning hyperparameters.

We again summarized all sweeps by calculating the “Usable Information”^17^ to quantify the direction and color information in RNNs, as shown in Fig. 5f and the results reaffirmed conclusions from the variance and decoding analyses. Together, these results suggest that a connection bottleneck in the form of neurophysiological architecture constraints (i.e., multiple areas, sparser connections between areas than within an area, as well as a smaller fraction of E-I connections) was the key design choice leading to RNNs with minimal color representations and consistent with the IB principle.

## Discussion

The goal of this study was to investigate if predictions from the IB principle in machine learning and information theory are also observed in cortical circuits. The IB principle defines an optimal representation to be one that retains only the relevant or useful information for solving a task^14^. This principle has been applied to explain the success of deep networks ^16,31^, by forming minimal sufficient representations of task inputs, leading to better generalization bounds and invariance to noise ^16^. We explored whether such a principle could explain cortical representations across different areas during a visual perceptual decision making task by probing the ability to decode task-relevant behavioral choices and external input information from different brain areas. We found that later areas of cortex along a sensorimotor transformation (in PMd) only represented the behavioral report, that is the action choice, while earlier areas had stronger input representations and performed relevant computations to define the behavioral report (in DLPFC). To better understand how such a phenomenon could be implemented in cortex, we trained many artificial multi-area RNNs to perform this task. Surprisingly, we also observed that RNNs formed minimal sufficient representations across a range of hyperparameter settings, suggesting the formation of minimal sufficient representations may be a more general feature of multi-area computation.

Given the “full-observability” of our multi-area models, we were able to analyze the learned weight matrices and understand how the network converged to transform task inputs into minimal sufficient representations by the output area. In particular, we found that the output-relevant direction information was preferentially propagated between areas by having the largest overlap with the top singular vector of the learned feedforward matrices. In contrast, color information was almost randomly propagated through feedforward connections. This mechanism is related to prior work on output potent and output null subspaces^32^ and communication subspaces ^1,33^, with the important difference that color information isn’t preferentially projected to a nullspace, but is aligned similarly to any random vector. Preferential alignment with a cortical nullspace is therefore not *necessary* to achieve an IB — color information may be attenuated through random alignment to a communication subspace. This solution (random alignment) poses less constraints on inter-areal connectivity than a solution that preferentially propagates direction information while also preferentially projecting color information to a nullspace.

Our results are also consistent with recent work proposing that cortical areas convey information through communication subspaces. One observation in communication subspaces is that they do not merely propagate the directions of highest variance^1^. We also observed this phenomenon for the **W**_21_ connectivity matrix, which communicates information from Area 1 to Area 2. Color activity had significant variance in Area 1 (see Fig. S8). Inter-areal connections must therefore not merely propagate the highest variance dimensions of a preceding area, otherwise color information would be conveyed to Area 2. Consistent with this, we found that while the top 2 PCs capture 97.7% excitatory unit variance, the top 2 readout dimensions of **W**_21_ only captured 40.0% of Area 1’s excitatory unit neural variance (Fig. S10). Hence, inter-areal connections are not aligned with the most variable dimensions, but are rather aligned to preferentially propagate certain types of information — a result consistent with a recent study analyzing links between activity in V1 and V2^1^.

We find minimal sufficient representations in PMd and in the later areas of our recurrent network models. Do such representations have any advantages? One possibility is that in cortex a minimal sufficient representation provides energetic benefits ^34,35^. Another possibility is that such a representation provides a computational advantage. This is an open question that is still somewhat unresolved in the machine learning community, with representation learning approaches that *maximize* mutual information between representations and inputs also leading to useful task representations^36^, in addition to compressed representations^16,17^. The information contained in the representation of a neural network is related to the “Information in the Weights” ^37^, which can be quantified using the Fisher Information ^38–40^, a measure of sensitivity to perturbations. This “Information in the Weights” view would predict that minimal sufficient representations have smaller Fisher information and are therefore less sensitive to (local) perturbations in the readout weights. In the context of deep networks, it has been proposed that minimal sufficient representations simplify the role of the output readout or classifier^16^. Further, a minimal sufficient representation with respect to a family of probabilistic decoders/classifiers will provably generalize better^41^.

Although finding a resolution to this debate in machine learning is beyond the scope of this paper, we assessed if minimal RNNs exhibited any qualities consistent with machine learning predictions. We explored whether minimal sufficient representations would simplify the readout, which we quantified by measuring the model’s performance in response to perturbations to the readout weights. We found that 3-area networks with minimal color information (particularly networks in Fig. 5b with no feedforward E-to-I connectivity) were less sensitive to perturbations than corresponding networks with significant color information (networks in Fig. 5a with 10% unconstrained feedforward connectivity, see Fig. S14). We also found that these networks differed significantly in readout complexity, with 3-area networks with minimal color information exhibiting simpler and sparser readouts (Fig. S14). However, we did not observe a clear trend between perturbation sensitivity and usable color information across random initializations (Fig. S14) for a fixed parameter setting (networks with 10% feedforward inhibition in Fig. 5b). An interesting venue for future work is to further examine the potential advantages of a minimal sufficient representation. Such findings would be valuable to the machine learning and neuroscience community. In our study, several factors including recurrent connectivity, multiple areas, and E/I populations make theoretical study of this question difficult. It is likely that studying this question requires simplifying the setting. For example, it likely makes sense to first focus on feedforward networks with a variable amount of task input information, similar to the generalized checkerboard-task used in a related study ^17^.

Our task could be solved with or without feedback connections with equivalent performance, indicating that feedback was not necessary to solve the task (Fig. S12). Minimal sufficient representations were found in both purely feedforward RNNs or RNNs with feedback (Fig. S12). When the model had feedback connections, we observed that feedback connections between Areas 2 and 1 preferentially conveyed direction information. Due to the presence of choice related signals in several cortical areas, these feedback connections may also play a role in computation of the direction choice. Another perspective on feedback signals is that they may related to error signals used for learning^42^. Multi-area networks may help understand and develop new hypotheses for physiological studies of feedforward and feedback computation ^43,44^, and more generally distributed processing for decision-making and cognition ^8,45^.Future research may use carefully designed tasks in conjunction with multi-area RNNs to better understand the role of feedback in computation.

We also found it was possible to solve this task with single area RNNs, although they did not resemble PMd (Figure S15) since it did not form a minimal sufficient representation. Rather, for our RNN simulations, we found that the following components were sufficient to induce minimal sufficient representations: RNNs with at least 3 areas, following Dale’s law (independent of the ratio of feedforward to feedback connections).

The multi-area RNN also provides several testable hypotheses. First, because the IB simplifies the readout, it suggests a simple readout from “output” areas of cortex (e.g PMd or Motor Cortex). In our multi-area RNN, we found that the output area was comprised of pools of neurons that represent left or right reaches^29^, enabling a simple readout (Fig. S13b, c.). In particular, the connectivity matrix for our area 3 was composed of two pools of excitatory neurons with a common inhibitory pool. Overall, this connectivity and winner-take-all architecture is consistent with our PMd activity, which appears to implement winner-take-all dynamics between a pool of neurons representing left and right reaches (Fig. S13h,i.) Second, due to selective propagation of inter-areal information, direction axis activity in DLPFC should be more predictive of activity in downstream regions such as PMdr and PMd than activity in the top PCs. Simultaneous recordings of DLPFC and PMd would help test this hypothesis.

## Materials and Methods

### Task and training details

#### Somatomotor reaction time visual discrimination task and recordings from DLPFC and PMd

The task, training and electrophysiological methods used to collect the data used here have been described previously ^23^ and are reviewed briefly below. All surgical and animal care procedures were performed in accordance with National Institutes of Health guidelines and were approved by the Stanford University Institutional Animal Care and Use Committee and the Boston University Institutional Animal Care and Use Committees.

Two trained monkeys (Ti and Ol) performed a visual reaction time discrimination task. The monkeys were trained to discriminate the dominant color in a central static checkerboard composed of red and green squares and report their decision with an arm movement. If the monkey correctly reached to and touched the target that matched the dominant color in the checkerboard, they were rewarded with a drop of juice. This task is a reaction time task, so that monkeys initiated their action as soon as they felt they had sufficient evidence to make a decision. On a trial-by-trial basis, we varied the signed color coherence of the checkerboard, defined as (*R* − *G*)*/*(*R* + *G*), where R is the number of red squares and G the number of green squares. The color coherence value for each trial was chosen uniformly at random from 14 different values arranged symmetrically from 90% red to 90% green. Reach targets were located to the left and right of the checkerboard. The target configuration (left red, right green; or left green, right red) was randomly selected on each trial. Both monkeys demonstrated qualitatively similar psychometric and reaction-time behavior. 996 units were selected from PMd of Ti (n=546) and Ol (n=450) and 2819 units were recorded from DLPFC of Ti while they performed the task^23^. Monkey Ol and Ti’s PMd units both had low choice color probability. Reported analyses from PMd data use units pooled across Monkey Ol and Ti.

#### RNN description and training

We trained a continuous-time RNN to perform the checkerboard task. The RNN is composed of *N* artificial neurons (or units) that receive input from *N*_in_ time-varying inputs **u**(*t*) and produce *N*_out_ time-varying outputs **z**(*t*). The RNN defines a network state, denoted by **x**(*t*) ∈ ℝ*^N^*; the *i*th element of **x**(*t*) is a scalar describing the “currents” of the *i*th artificial neuron. The network state is transformed into the artificial neuron firing rates (or network rates) through the transformation:

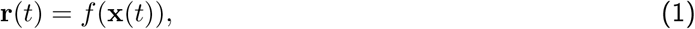

where *f* (·) is an activation function applied elementwise to **x**(*t*). The activation function is typically nonlinear, endowing the RNN with nonlinear dynamics and expressive modeling capacity ^46^. In this work, we use *f* (*x*) = max(*x,* 0), also known as the rectified linear unit, i.e., *f* (*x*) = relu(*x*). In the absence of noise, the continuous time RNN is described by the equation

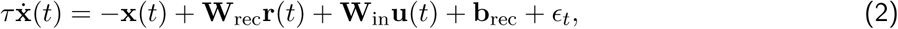

where *τ* is a time-constant of the network, **W**_rec_ ∈ ℝ*^N^*^×^*^N^* defines how the artificial neurons are recurrently connected, **b**_rec_ ∈ ℝ*^N^* defines a constant bias, 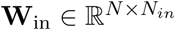 maps the RNN’s inputs onto each artificial neuron, and *E_t_* is the recurrent noise. The output of the network is given by a linear readout of the network rates, i.e.,

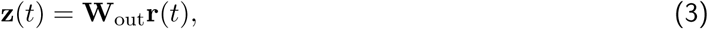

where 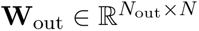 maps the network rates onto the network outputs.

We trained RNNs to perform the checkerboard task as follows. For all networks, unless we explicitly varied the amount of units, we used *N*_in_ = 4, *N* = 300, and *N*_out_ = 2.

The four inputs were defined as:

1. Whether the left target is red (−1) or green (+1).
2. Whether the right target is red (−1) or green (+1).
3. Signed coherence of red (ranging from −1 to 1), (*R* − *G*)*/*(*R* + *G*).
4. Signed coherence of green (ranging from −1 to 1), (*G* − *R*)*/*(*R* + *G*). Note that, prior to the addition of noise, the sum of the signed coherence of red and green is zero.

The inputs, **u**(*t*) ∈ ℝ^4^, were defined at each time step, *t*, in distinct epochs. In the ‘Center Hold’ epoch, which lasted for a time drawn from distribution 𝒩 (200 ms, 50^2^ ms^2^), all inputs were set to zero. Subsequently, during the ‘Targets’ epoch, which lasted for a time drawn from distribution 𝒰[600 ms, 1000 ms], the colors of the left and right target were input to the network. These inputs were noiseless, as illustrated in Fig. 3a, to reflect that target information is typically unambiguous in our experiment. Following the ‘Targets’ epoch, the signed red and green coherences were input into the network during the ‘Decision’ epoch. This epoch lasted for 1500 ms. We added zero mean independent Gaussian noise to these inputs, with standard deviation equal to 5% of the range of the input, i.e., the noise was drawn from 𝒩 (0, 0.1^2^). At every time point, we drew independent noise samples and added the noise to the signed red and green coherence inputs. We added recurrent noise *E_t_*, adding noise to each recurrent unit at every time point, from a distribution 𝒩 (0, 0.05^2^). Following the ‘Decision’ epoch, there was a ‘Stimulus Off’ epoch, where the inputs were all turned to 0.

The two outputs, **z**(*t*) ∈ ℝ^2^ were defined as:

1. Decision variable for a left reach.
2. Decision variable for a right reach.

We defined a desired output, **z**_des_(*t*), which was 0 in the ‘Center Hold’ and ‘Targets’ epochs. During the ‘Decision’ epoch, **z**_des_(*t*) = 1. In the ‘Stimulus Off’ epoch, **z**_des_(*t*) = 0. In RNN training, we penalized output reconstruction using a mean-squared error loss,

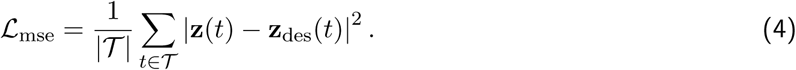

The set 𝒯 included all times from all epochs except for the first 200 ms of the ‘Decision’ epoch from the loss. We excluded this time to avoid penalizing the output for not immediately changing its value (i.e., stepping from 0 to 1) in the ‘Decision’ epoch. Decision variables are believed to reflect a gradual process consistent with non-instantaneous integration of evidence, e.g., as in drift-diffusion style models, rather than one that steps immediately to a given output.

To train the RNN, we minimized the loss function:

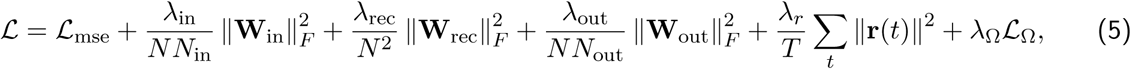

where

■ ||**A**||*_F_* denotes the Frobenius norm of matrix **A**.
■ *λ*_in_ = *λ*_rec_ = *λ*_out_ = 1*, λ_r_* = 0 to penalize larger weights.
■ *λ*_Ω_ = 2.
■ ℒ_Ω_ is a regularization term that ameliorates vanishing gradients proposed and is described in prior literature^26,47^.

During the training process, we also incorporated gradient clipping to prevent exploding gradients^47^. Training was performed using stochastic gradient descent, with gradients calculated using backpropagation through time. For gradient descent, we used the Adam optimizer, which is a first order optimizer incorporating adaptive gradients and momentum^48^.

Every 200 or 500 training epochs, we generated 2800 cross-validation trials, 100 for each of the 28 possible conditions (14 coherences × 2 target configurations). For each trial, there was a correct response (left or right) based on the target configuration and checkerboard coherence. When training, we defined a “correct decision” to be when the RNNs DV for the correct response was greater than the other DV and the larger DV was greater than a pre-set threshold of 0.6. We evaluated the network 500ms before the checkerboard was turned off (the end of the trial). We required this criteria to be satisfied for at least 65% of both leftward and rightward trials. We note that this only affected how we terminated training. It had no effect on the backpropagated gradients, which depended on the mean-squared-error loss function. Note that a trial that outputted the correct target but did not reach the 0.6 threshold would not be counted towards the 65% criteria.

When testing, we defined the RNNs decision to be either: (1) whichever DV output (for left or right) first crossed a pre-set threshold of 0.6, or (2) if no DV output crossed the pre-set threshold of 0.6 by the end of the ‘Decision epoch,’ then the decision was for whichever DV had a higher value at the end of this epoch — an approach that is well established in models of decision-making ^49,50^. If the RNN’s decision on a single trial was the same as the correct response, we labeled this trial ‘correct.’ Otherwise, it was incorrect. The proportion of decisions determined under criterion (2) was negligible (0.5% across 100 trials for each of 28 conditions). An interpretation for criterion (2) is that if the RNN’s DV has not achieved the threshold certainty level by the end of a trial, we assign the RNN’s decision to be the direction for which its DV had the largest value. Finally, in training only, we introduced ‘catch’ trials 10% of the time. On 50% of catch trials, no inputs were shown to the RNN and **z**_des_(*t*) = 0 for all *t*. On the remaining 50% of catch trials, the targets were shown to the RNN, but no coherence information was shown; likewise, **z**_des_(*t*) = 0 for all *t* on these catch trials.

We trained the three-area RNNs by constraining the recurrent weight matrix **W**_rec_ to have connections between the first and second areas and the second and third areas. In a multi-area network with *N* neurons and *m* areas, each area had *N/m* neurons. In our 3-area networks, each area had 100 units. Of these 100 units, 80 were excitatory and 20 were inhibitory. Excitatory units were constrained to have only positive outgoing weights, while inhibitory units were constrained to have only negative outgoing weights. We used the pycog repository ^26^ to implement these architecture constraints. The parameters for the exemplar RNN used in the paper are shown in Table 1. In our hyperparameter sweeps, we varied the hyperparameters of the exemplar RNN. For each parameter configuration, we trained 8 different networks with different random number generator seeds. For analyses of the RNN, we fixed the timing of trials, obviating the need to to restretch trial lengths. Note that while at inference, we generated RNN trials with equal length, the RNN was trained with varying delay periods.

**Table 1:**
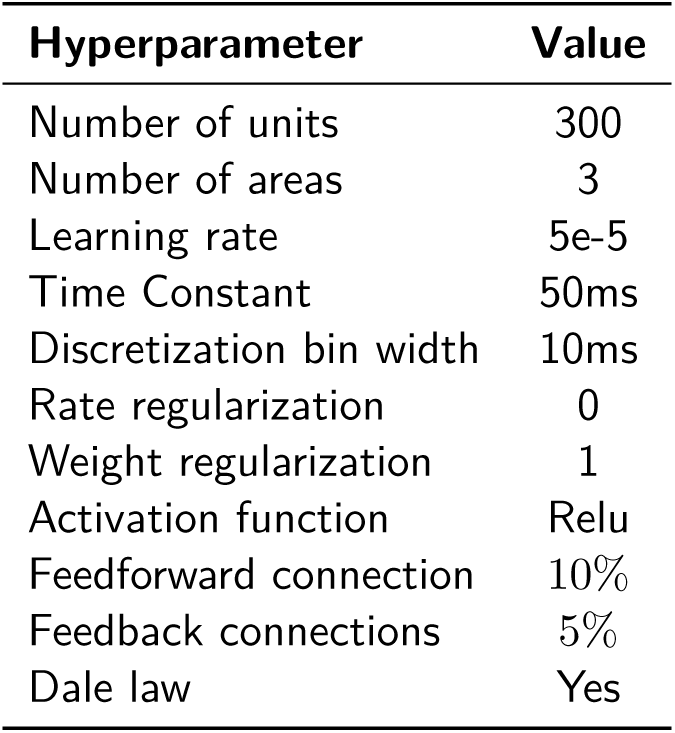
Hyperparameters of exemplar RNN.

| Hyperparameter | Value |
| --- | --- |
| Number of units | 300 |
| Number of areas | 3 |
| Learning rate | 5e-5 |
| Time Constant | 50ms |
| Discretization bin width | 10ms |
| Rate regularization | 0 |
| Weight regularization | 1 |
| Activation function | Relu |
| Feedforward connection | 10% |
| Feedback connections | 5% |
| Dale law | Yes |

### Additional description of analyses

#### Decoding analysis for DLPFC and PMd data

For DLPFC and PMd data, we calculated decoding accuracy using 400 ms bins. We report numbers in a window [−300ms, +100 ms] aligned to movement onset. We used the Python *sklearn.svm.SVC* command with 80% training and 20 % test sets. Decoding analyses were performed on single trials using 2-49 simultaneously recorded units from Plexon U-probes and the averages reported are across PMd and DLPFC sessions, respectively. To assess whether decoding accuracies were significant, we shuffled the trials for 100 times. The decoding accuracy for direction, color and target configuration variables of a session was judged to be significant if it lies above the 99 percentile of shuffled accuracy. For DLPFC, direction, target configuration and color decoding accuracy of 100%, 78%, 53% sessions were judged to be significantly above chance; for PMd, 97%, 26%, 20% sessions demonstrated significant decoding accuracy to direction, target configuration and color.

Mutual information was calculated by computing *H*(*Y*) − *L_CE_* where *H*(*Y*) was 1 and *L_CE_* denotes the cross entropy loss (in bits). We computed the decoding and information only for sessions with decoding accuracy significantly above chance. Negative mutual information was set to zero.

#### Decoding and Mutual information for RNNs

We used a decoder and mutual information approximation to quantify the amount of information (color, target configuration, direction) present in the network. In particular, we trained a neural network to predict the reported direction choice, color choice, and target configuration from the single trial activity of a population of units. We focused on decoding the network’s choice (as described above in *“RNN description and training”* of Methods) rather than the ground-truth input information to focus on how the behavioral choice was represented across areas. We note that the network’s choice and input information are highly correlated in trained networks (Fig. 1b and Fig. S1). We used 700 trials for training, and 2100 independent trials for testing. To generate the trials for training and testing, we increased the recurrent noise to be drawn from the distribution (𝒩 (0, 0.1^2^)) to prevent overfitting. For each trial, we averaged data in a window [−300ms, +100ms] around reaction time.

We trained a neural network with 3 layers, 64 units per layer, leakyRelu activation (*α*=0.2), and dropout (p=0.5), using stochastic gradient descent, to predict the choice given the activity of the population. We trained the neural network to minimize the cross-entropy loss. We used the same neural network from the decode to compute an approximation to mutual information, described in Supplementary Note 2. In previous work^29^, we obtained analogous results using a linear decoder across hyperparameter sweeps. We also obtain analogous results by using an SVM decoder in Fig. S4 for our exemplar parameter configuration.

#### RNN behavior

To evaluate the RNN’s psychometric curve and reaction-time behavior, we generated 200 trials for each of the 28 conditions, producing 400 trials for each signed coherence. For these trials, we calculated the proportion of red decisions by the RNN. This corresponds to all trials where the DV output for the red target first crossed the preset threshold of 0.6; or, if no DV output crossed the threshold of 0.6, if the DV corresponding to the red target exceeded that corresponding to the green target. The reaction time was defined to be the time between checkerboard onset to the first time a DV output exceeded the preset threshold of 0.6. If the DV output never exceeded a threshold of 0.6, in the reported results, we did not calculate a RT for this trial.

#### dPCA

Demixed principal components analysis (dPCA) is a dimensionality reduction technique that provides a projection of the data onto task related dimensions while preserving overall variance^51^. dPCA achieves these aims by minimizing a loss function:

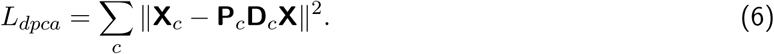

Here, **X***_c_* refers to data averaged over a “dPCA condition” (such as time, coherence, target configuration, color, or direction), having the same shape as **X** ∈ ℝ*^N^*^×^*^cT^*, but with the entries replaced with the condition-averaged response. The aim is to recover (per dPCA condition *c*) a **P***_c_* and **D***_c_* matrix. **P***_c_* is constrained to have orthonormal columns, while **D***_c_* is unconstrained. The number of columns of **P***_c_* and rows of **D***_c_* reflects the number of components one seeks to find per condition. The column of **P***_c_* reflects how much the demixed data contributes to each neuron. We use the principal axes from **P***_c_* to compute the axis overlap, as in Kobak et al^51^. We used axes of dimension 1 for RNNs, which were sufficient to capture most color, target configuration, or direction variance. For the neural data, we used five components for direction, color and target configuration since the PMd data was higher dimensional than the RNNs.

For multi-area analyses, we separated the units for each area and found the task-relevant axes for this subset of units. For the inter-area analyses, we used RNNs with only excitatory connections, and therefore found the color and direction axis using only the excitatory units. In all other analyses, all units were used to identify the axes. For RNN activity, we performed dPCA using activity over the entire trial. For analyses of the RNN, we fixed the timing of trials, obviating the need to to restretch trial lengths. Note that while at inference, we generated RNN trials with equal length, the RNN was trained with varying delay periods.

For neural data, due to the stochasticity in task design, there is a trial-by-trial difference in interval between target and checkerboard onset (TC interval). The reaction time (from the checkerboard onset to monkey’s hand movement initiation) also varies for each trial. To align time events across trials, we restretched the firing rates in each trial. For DLPFC units, each trial was aligned to targets onset first. Median reaction time (527ms) and TC interval (735ms) were calculated by combining every trial in the database. For each trial, TC interval and reaction time was either compressed or stretched to the median values through linear interpolation. After the data restretching, we choose the data window **T** as 1300ms, from −100ms to 1200ms around target onset with sample size of 1ms. For every unit **n** in total units number **N**, we averaged the single-trial firing rate by stimulus **S** (checkerboard dominant color, green or red) and decision of choice **D** (left or right). As a result, a 4D firing-rate matrix **X^N^**^×^**^S^**^×^**^D^**^×^**^T^** was created as input to demixed principal component analysis algorithm.

For PMd units, activities before checkerboard onset were minimal. As a result, each trial was aligned to target onset and a segment with time window of [−100*ms,* 367*ms*] was chosen first. Then the same trial was aligned to checkerboard first and a segment with a window of [−368*ms,* 465*ms*] was chosen. The final restretched data was the concatenation of these two data segments.

When computing the overlap in Fig. 4, we averaged across 8 initializations, and computed the PSTHs over 700 trials. In our dpca variance sweeps (Fig. 5), we computed the PSTHs over 280 trials.

#### PCA

Principal components analysis (PCA) is a dimensionality reduction technique that projects high-dimension data into low-dimensional axis which maximize the variance in the data. PCA provides low-dimensional projections by minimizing the loss function:

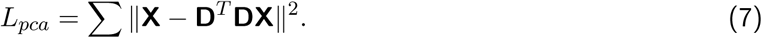

**X^N^**^×^**^T^** is high-dimension raw data and **D^N^**^×^**^N^** is the decoding matrix. The low-dimension trajectories **x^M^**^×^**^T^** (*M < N*) calculated by multiplying first *M* rows of **D** by **X**.

Before applying PCA on the data, the raw data was preprocessed by data normalization and average firing rate removal:

#### Data normalization for PCA for neural data

Due to the hererogenity in firing rates, PCA results might be dominanted by units with high firing rates. To equalize the contribution of each unit in PCA, for each unit **n** ∈ 𝒩, 99 percentile of max firing rate **x_n,99_** was calculated over all stimulus, decision conditions and time point. Then we divided each data for unit **x_n,99_** by square root of to normalize the data.

#### Condition independent signal removal for PCA

The condition independent signal is another source that explains substantial amount of population variance other than task-related signal. Before conducting principal component analysis (PCA), we calculated the average firing rate of each single unit **X**^1×1×1×^**^T^** over all stimulus and decision conditions and subtracted this condition independent signal from the time-restretched data **X**^1×^**^S^**^×^**^D^**^×^**^T^**. We also visualized the PCs of the RNNs in an analogous manner, unless otherwise noted.

#### Canonical correlation

We applied CCA to assess the similarity between neural activity and the artificial unit activity ^52^. Before applying CCA, we performed principal component analysis to reduce the dimensionality of the artificial and neural activity to remove noise, which can be arbitrarily reshaped to increase the canonical correlation ^52^. We reduced the dimensionality of PMd data to 2 (which captures over 80% of the PMd variance). For DLPFC, 18 dimensions were required to capture over 80% of the variance, but at such a high dimensionality, noise can be reshaped to significantly increase the canonical correlations. For DLPFC, we therefore show a comparison to the top 4 PCs in Fig. 3e. However, the trends held irrespective of the DLPFC dimensionality we chose, as shown in Fig. S3. We report the average CCA correlation coefficients in Fig. 3e using times in a window of [−400ms, 400ms] aligned to checkerboard onset. The data was binned in 10ms bins.

#### Analyses of inputs and activity

In order to disentangle the effects of external inputs and recurrence, in Fig. S9, we evaluated the input contribution and overall activity. For Area 1, we defined the input contribution as **W**_in_**u***_t_*, and for areas 2 and 3, we defined the input contribution as 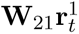, and 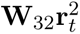 respectively, where 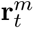 denotes the activity of the units in area *m*. The activity 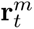 corresponds to the firing rate that experimentalists could measure, reflecting a combination of input and recurrent interactions. For constant inputs, a stable value of the activity implies there is little recurrent processing.

#### Inter-Area Projection Analyses

To calculate the overlap between the color and direction axes with the potent and null spaces, we performed singular value decomposition on the inter-area connections, **W**_21_ and **W**_32_. **W**_21_ and **W**_32_ were 80 × 80 matrices, and were full rank. Nevertheless, they had near some zero singular values, indicating that the effective rank of the matrix was less than 80. We defined the potent dimensions to be the top *m* right singular vectors, while the null dimensions were the remaining 80 − *m* right singular vectors.

We performed the analyses of Fig. 4f by varying the potent and null dimensions, sweeping *m* from 1 to 80. For each defined potent and null space, we calculated the axis overlap between the direction (or color) axis and the potent (or null) space by computing the L2-norm of the orthogonal projection (squared). We report the squared quantity because the expectation of the norm of a projection of a random vector onto an *m*-dimensional subspace of an *n*-dimensional space is *m/n*. We include an approximation of the expectation of the projection of a random vector in Fig. 4 by averaging the projection of 100 random vectors. Our results show that the direction axis was always more aligned with potent dimensions than the color axis, irrespective of the choice of *m*, and that the direction axis was preferentially aligned with the top singular vector.

## Supplementary Information for: “A cortical information bottleneck during decision-making”

**Figure S1:**
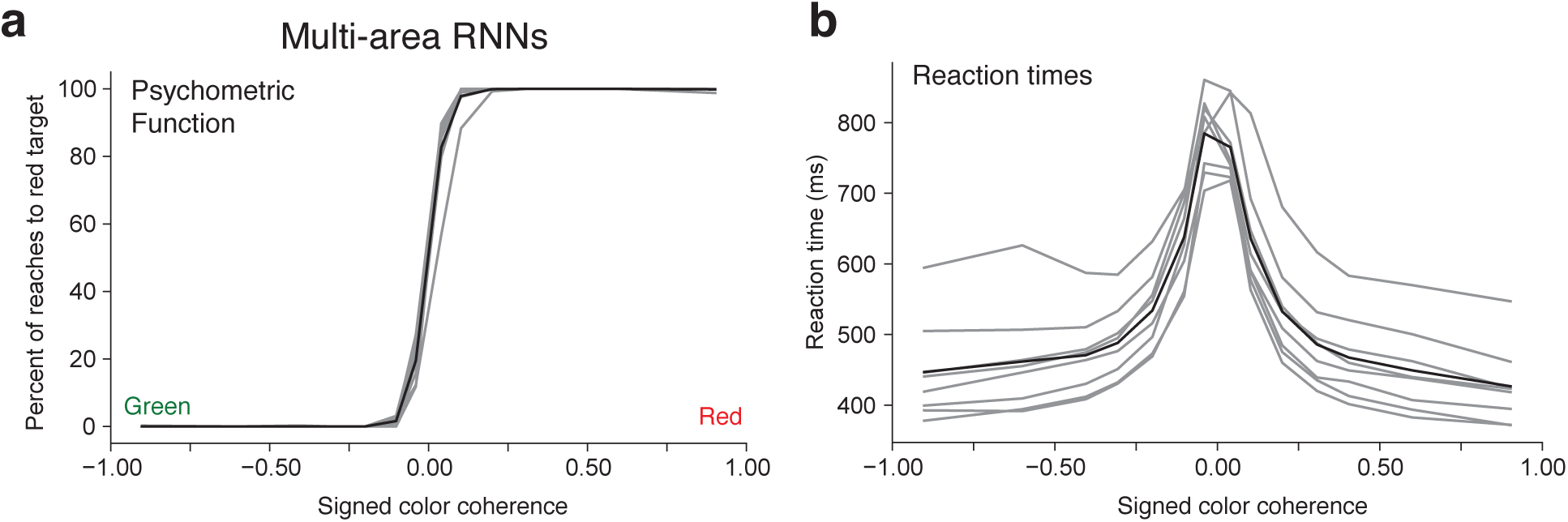
**(a)** Psychometric and **(b)** reaction time curves for multi-area RNNs. The hyperparameters used for these RNNs are described in Table 1. Gray lines represent individual RNNs and the black solid line is the average across all RNNs.

**Figure S2:**
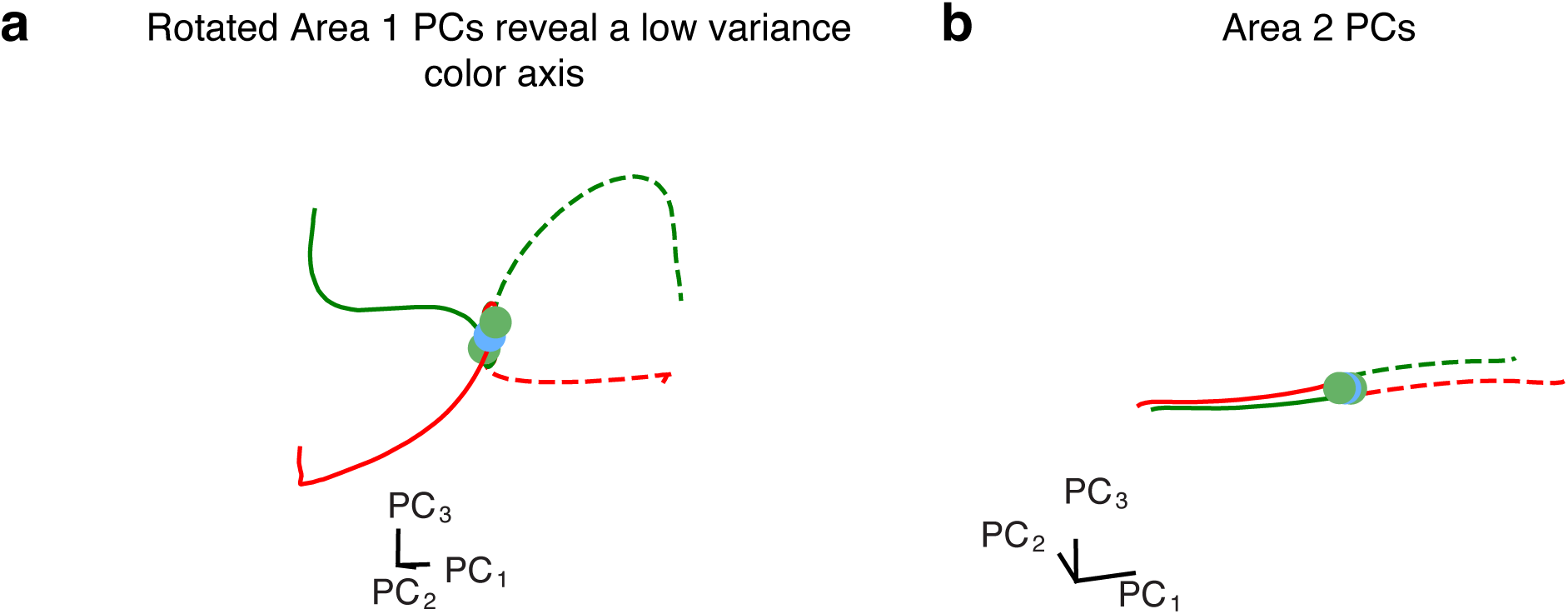
**(a)** Another rotation of the first three PCs for Area 1 RNN, with PC_3_ amplified to show that there is a low variance color axis. **(b)** Area 2 PCs in the same projection as used in Figure 3. While these PCs qualitatively appear to represent the direction decision, they are distinct from Area 3, with Area 3 demonstrating a stronger resemblance to PMd activity.

**Figure S3:**
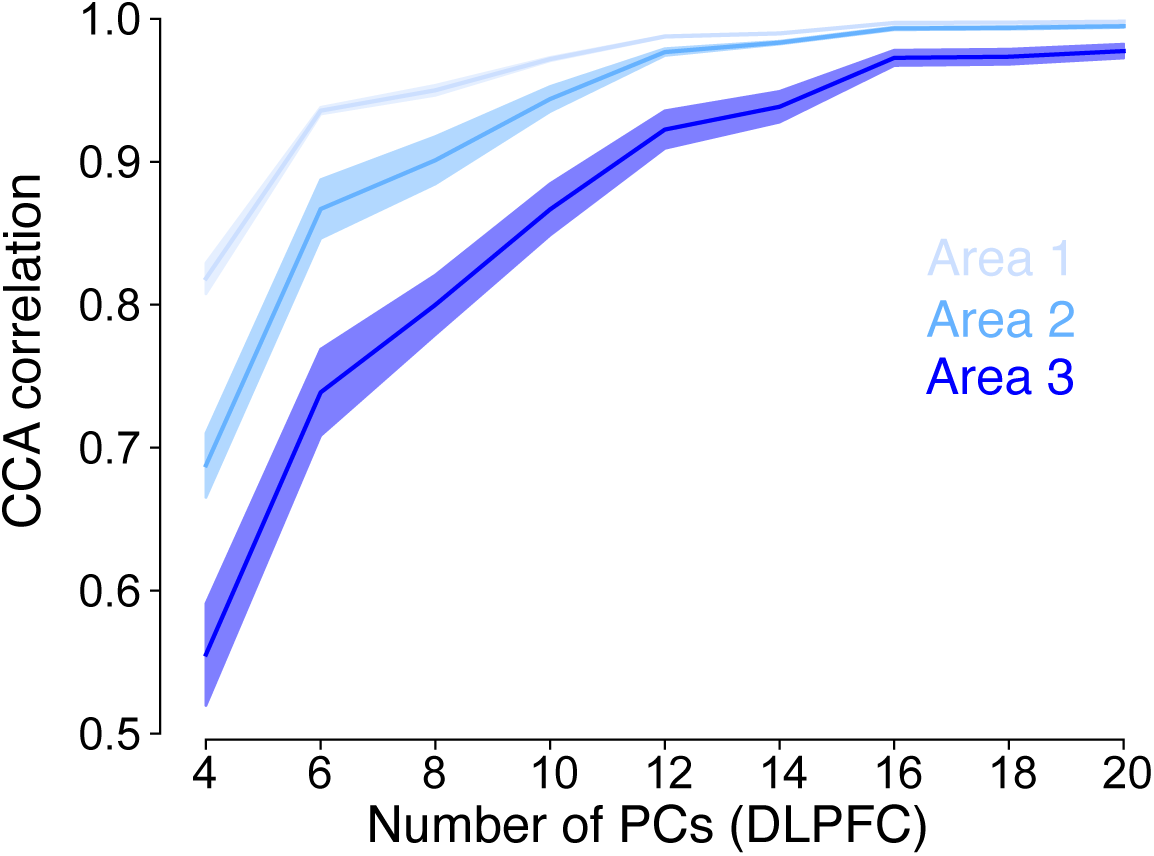
Because DLPFC is higher-dimensional than PMd, we performed the CCA correlation coefficient comparison to Areas 1-3 of the RNN varying the number of dimensions used for the DLPFC PCs. Note that as dimensionality increases, CCA correlation coefficient increases because additional dimensions, which are low variance, can be weighted to better reproduce the RNN PCs. We nevertheless observe that Area 1 has the highest CCA correlation to DLPFC, while Area 3 has the least.

**Figure S4:**
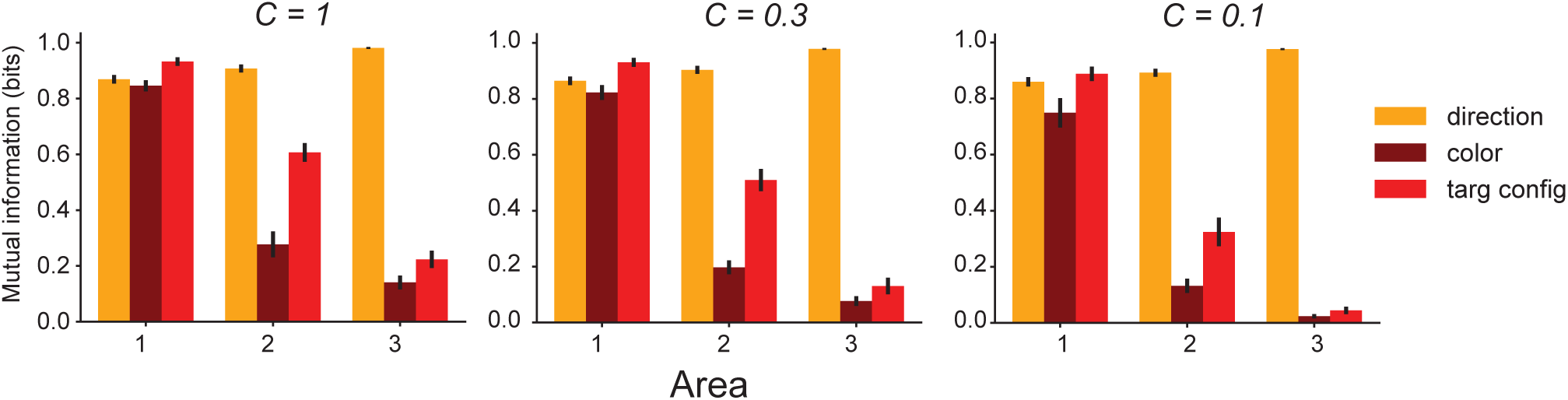
SVM Mutual Information (approximated using the Usable Information) for each RNN area as a function of increasing decoder L2 regularization *C*. A lower *C* implies more regularization.

**Figure S5:**
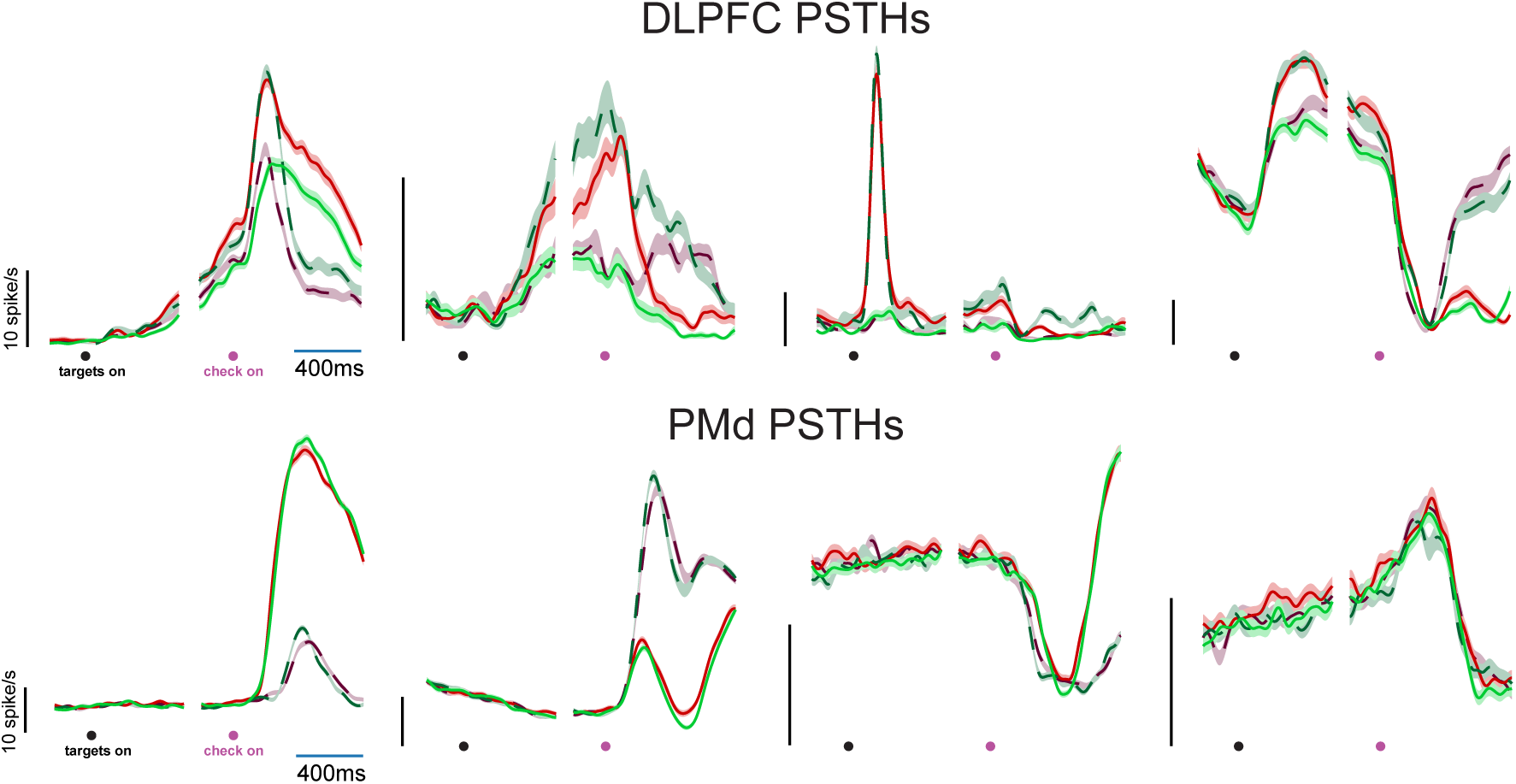
DLPFC and PMd units aligned to targets and checkerboard with a breakout in the middle due to trial-by-trial difference in interval between these two stimuli.

**Figure S6:**
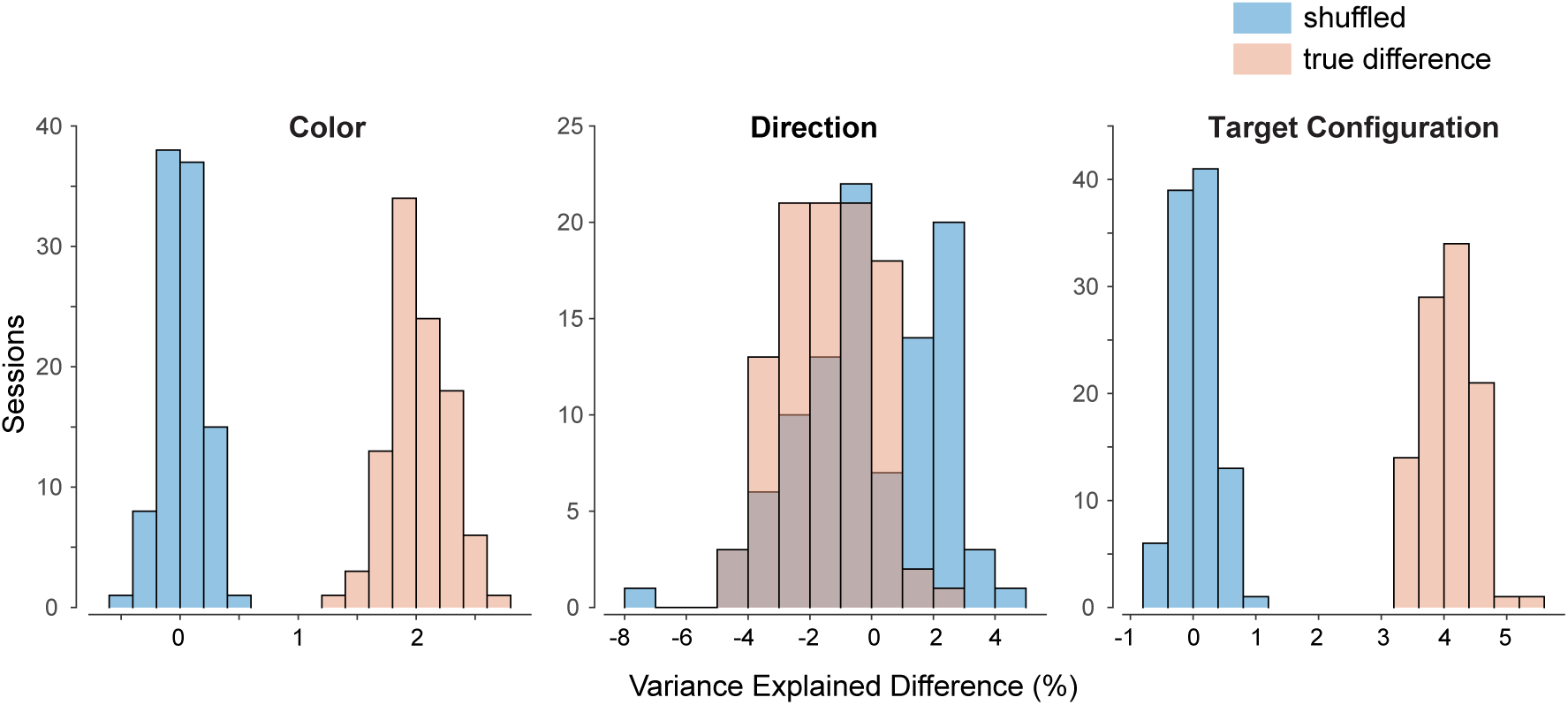
Shuffle test for assessing significance of dPCA results. To assess whether the dPCA explained variances for DLPFC and PMd are significantly different from one another, we performed the following analysis. We randomly sampled 500 units from the DLPFC dataset and 500 units from the PMd dataset. We then used dPCA to measure the variance explained by target configuration, color choice, and reach direction. We combined the PMd and DLPFC dataset into a pool of 1000 units and then randomly selected 500 units from this pool to create a surrogate PMd dataset and used the remaining 500 units as a surrogate DLPFC dataset. We again performed dPCA on these surrogate datasets and estimated the variance for task variables. We repeated this process for 100 times and estimated a sampling distribution for the true difference in variance between DLPFC and PMd for various task variables. At the same time, we estimated the distribution of the variance difference between surrogate PMd and DLPFC dataset for various task variables. We then defined a p-value as the number of shuffles in which the difference in variance was higher than the median of the true difference and divided it by 100. The differences were statistically significant (p < 0.02) for color and target configuration but not for direction (p=0.72).

**Figure S7:**
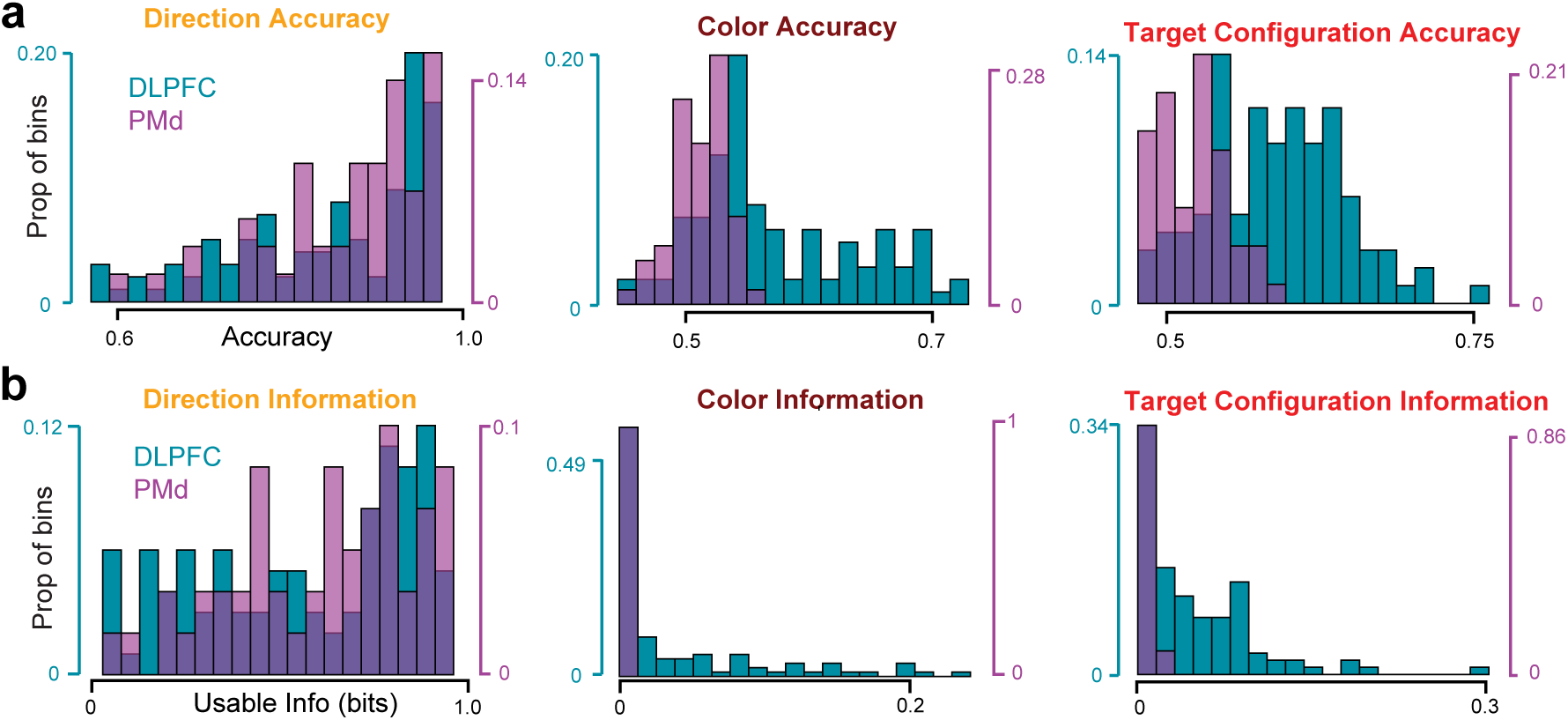
Histogram (across sessions) of direction, color, and target configuration decode accuracy **(a)** and **(b)** usable information for DLPFC and PMd of all sessions (including those sessions with decoding accuracy below 0.5).

**Figure S8:**
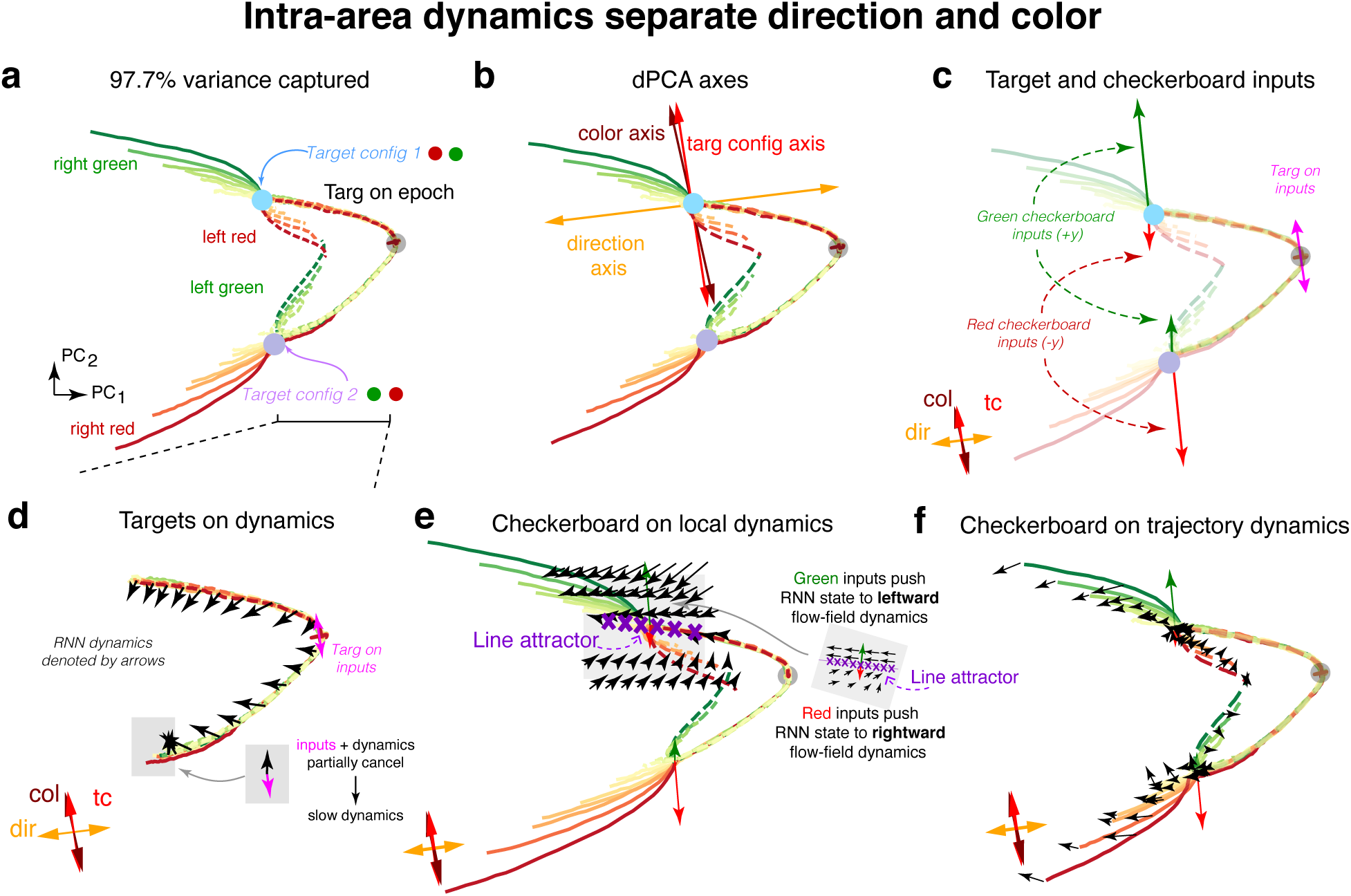
Candidate mechanism for axis orthogonalization. **(a)** Top 2 PCs of RNN Area 1 activity. Trajectories are now colored based on the coherence of the checkerboard, and the condition-independent signal is not removed. We did not remove the condition-independent signal so we could directly study the high-dimensional dynamics of the RNN and its equilibrium states. The trajectories separate to two regions corresponding to the two potential target configurations (Target config 1 in blue, Target config 2 in purple). The trajectories then separate upon checkerboard color input, leading to four trajectory motifs. **(b)** Projection of the dPCA principal axes onto the PCs. **(c)** Projection of the target configuration and color inputs onto the PCs. Target configuration inputs are shown in pink, a strongly green checkerboard in green, and a strongly red checkerboard in red. Irrespective of the target configuration, green checkerboards cause the RNN state to increase along PC_2_ while red checkerboards cause the RNN state to decrease along PC_2_. The strength of the input representation is state-dependent: checkerboards corresponding to left reaches, whether they are green or red, cause smaller movements of the RNN state along the color axis. **(d)** Visualization of RNN dynamics and inputs during the target presentation. In the Targets On epoch, target configuration inputs cause movement along the vertical target configuration axis. The RNN dynamics implemented a leftward flow-field that pushed the RNN state into an attractor region of slow dynamics. **(e)** At the Target config 1 attractor, we plot the local dynamics using a previously described technique ^53^. The RNN implements approximately opposing flow fields above and below a line attractor. The line attractor, denoted by purple crosses, is characterized by slow dynamics. Above the attractor, a leftward flow-field increases direction axis activity, while below the attractor, a rightward flow-field decreases direction axis activity. A green checkerboard input therefore pushes the RNN state into the leftward flow-field (solid green trajectories) while a red checkerboard input pushes the RNN state into a rightward flow-field (dotted red trajectories). This computes the direction choice in a given target configuration, while allowing the direction axis to be orthogonal to color inputs. Arrows are not to scale; checkerboard inputs have been amplified to be visible. **(f)** Visualized dynamics across multiple trajectory motifs. These dynamics hold in both target configurations leading to separation of right and left decisions on the direction axis. Arrows are not to scale, for visualization purposes.

**Figure S9:**
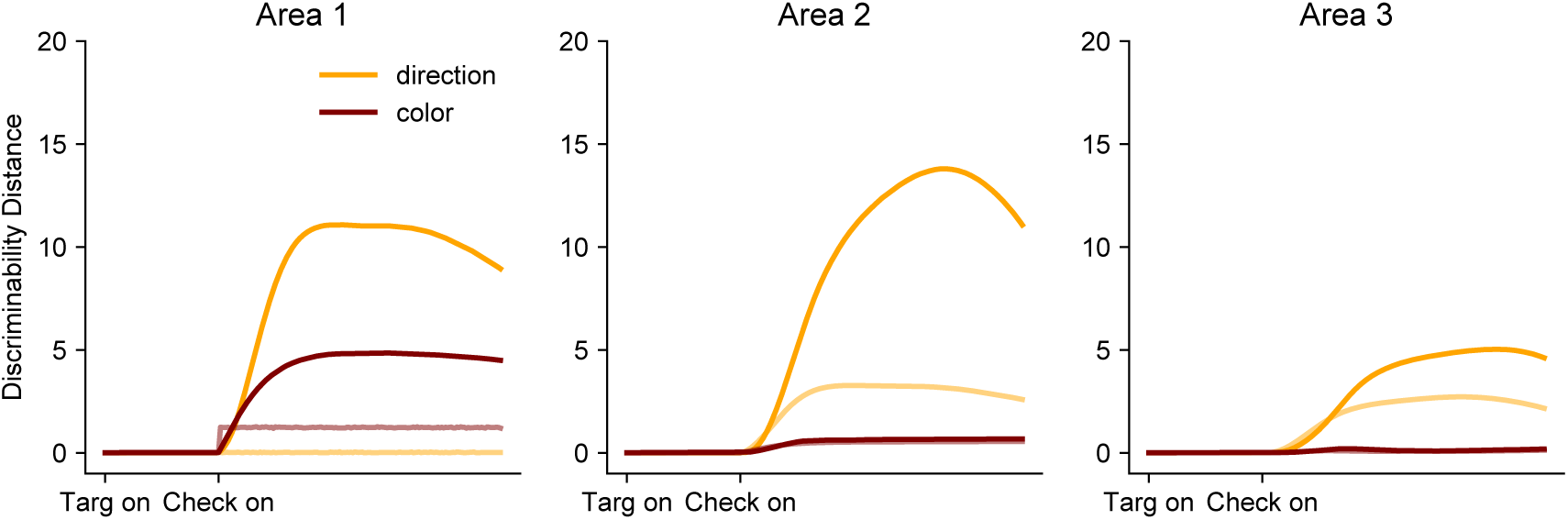
The norm of the direction discriminability (left red - right red + left green - right green)/2 and color discriminability (left green - left red + right green - right red)/2 as a function of the processing area. The inputs are shown in lighter transparency and the overall activity is shown in solid lines. Area 1 has significant recurrence evidenced by a large separation between the input and overall activity. For our exemplar network, there is very little evidence of recurrent filtering of color information (i.e recurrent activity is never below inputs).

**Figure S10:**
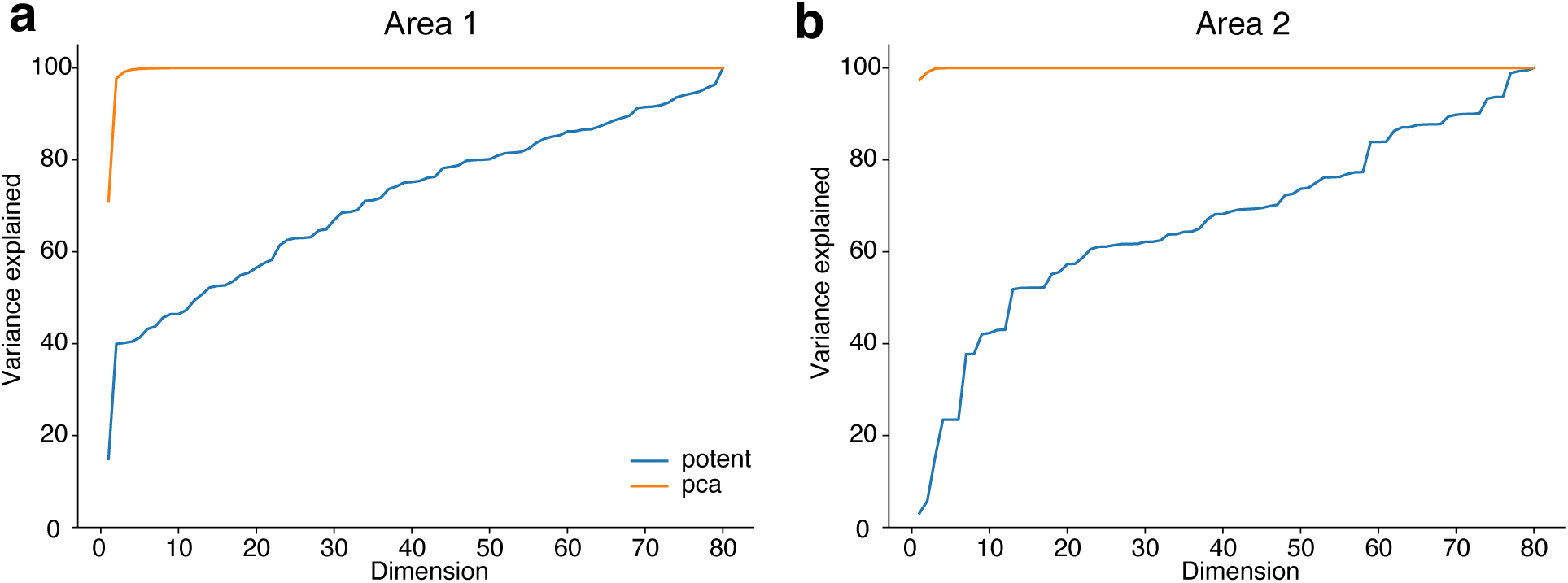
Relationship between PCs and inter-area potent space. **(a)** Variance explained of the excitatory units in Area 1 by the top principal components and top dimensions of potent space of **W**_21_, swept across all dimensions. **(b)** Variance explained of the excitatory units in Area 2 by the top principal components and top dimensions of potent space of **W**_32_, swept across all dimensions. These plots show that the connections between areas do not necessarily propagate the most dominant axes of variability in the source area to the downstream area. Excitatory units were used for the comparison because only excitatory units are read out by subsequent areas. These results were upheld when comparing to the variance explained by the top principal components obtained from all units.

**Figure S11:**
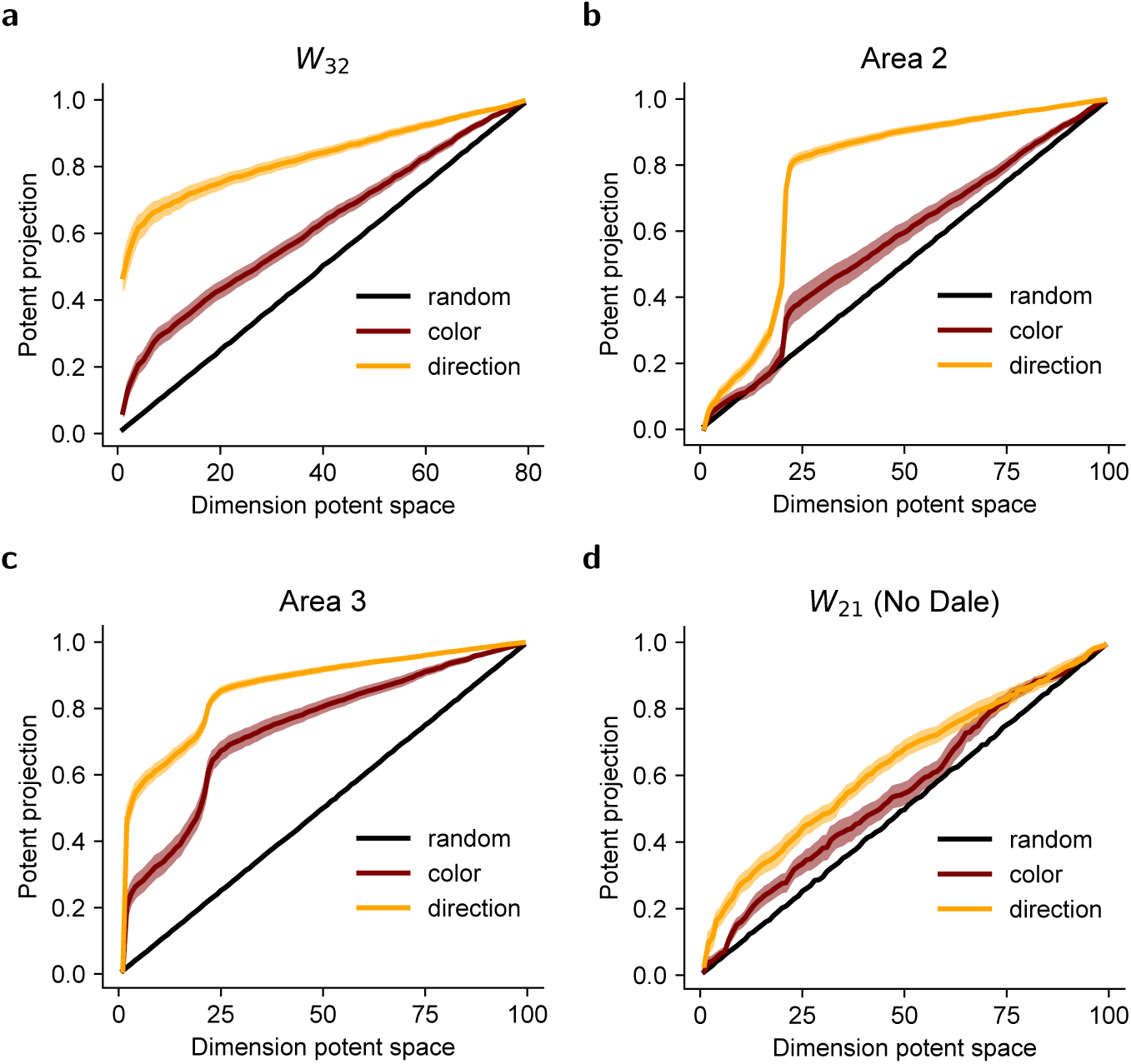
**(a)** Alignment of dPCA color and direction axes from area 2 with inter-areal connections **W**_32_. **(b,c)** Alignment of dPCA axes with intra-areal recurrent matrices for 3 area dale networks (Area 2 and Area 3). **(d).** Alignment of dPCA axes in area 1 with **W**_21_ for networks without Dale’s law. In contrast to Fig. 4f, direction information is not preferentially propagated. Same conventions as Fig. 4c,f.

**Figure S12:**
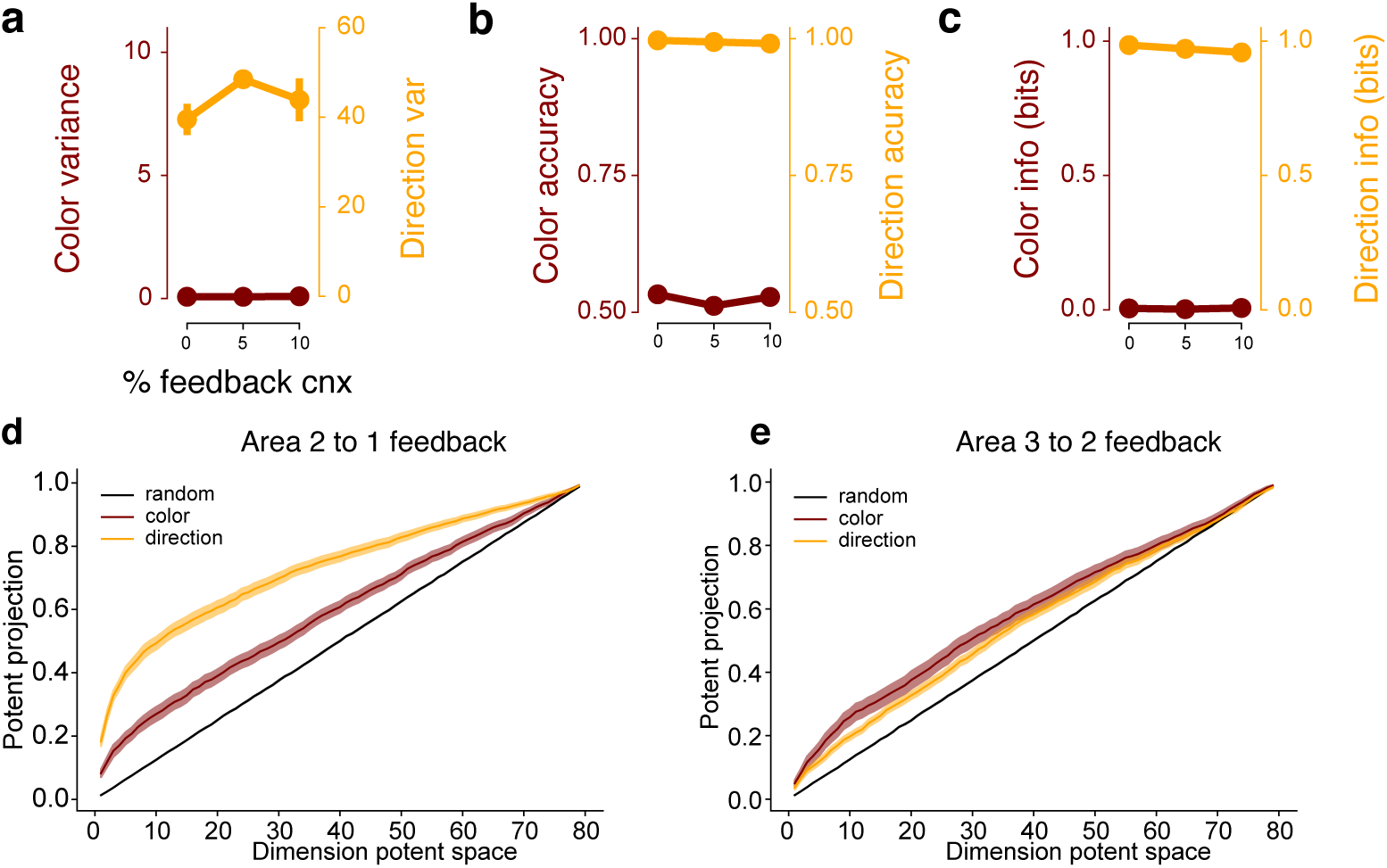
Effect of feedback connections. **(a)** dPCA variance in area 3 of RNNs where we varied the amount of feedback connectivity. RNNs exhibited nearly zero dPCA color variance in Area 3 across networks with 0%, 5%, and 10% feedback connections. **(b, c)** RNNs also exhibited minimal color representations, achieving nearly chance levels of decode accuracy and nearly zero mutual information. **(d, e)** Feedback projections of the color and direction axis on the feedback inter-area matrix between **(d)** area 2 and area 1, and **(e)** area 3 and area 2 (for networks trained with 5% feedback connections, across variable feedforward connectivity percentages).

**Figure S13:**
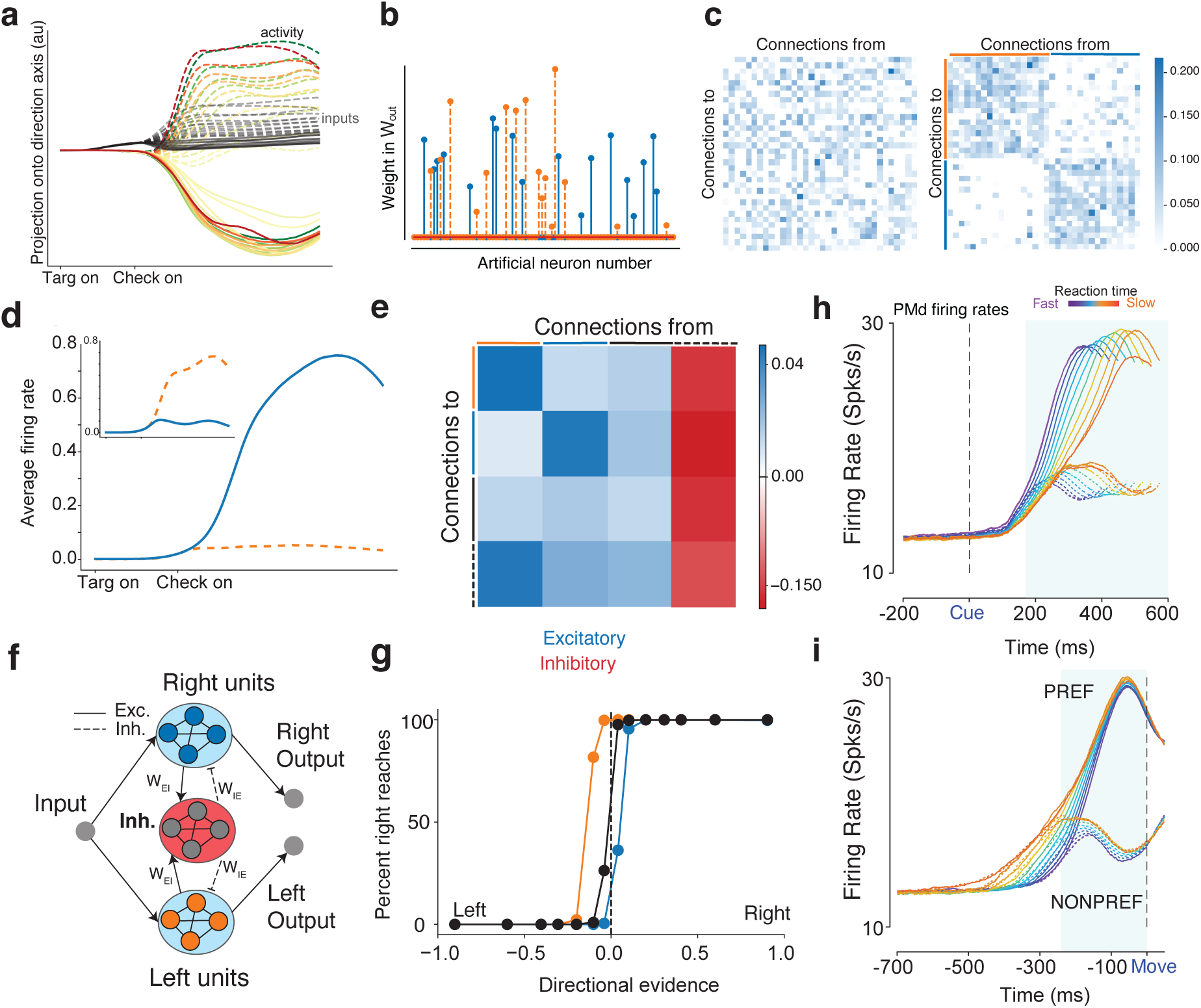
Area 3 mechanism. **(a)** Projection of input and overall activity onto the direction axis identified through dPCA. **(b)** Readout weights in **W**_out_ are sparse, with many zero entries, and selective weights for a left or right reach. **(c)** The unsorted connectivity matrix for the nonzero readout units (left panel), and the sorted connectivity matrix when the matrix was reordered based on the readout weight pools (right). **(d)** Average PSTHs from units for a leftward reach and (inset) rightwards reach. When one pool increases activity, the other pool decreases activity. **(e)** Averaged recurrent connectivity matrix. **(f)** Schematic of output area. **(g)** Psychometric curve after perturbation experiment, where 10% of inhibitory weights to the left pool (orange) and right pool (blue) were increased (doubled). Directional evidence is computed by using the signed coherence and using target configuration to identify the strength of evidence for a left reach and strength of evidence for a right reach. Increasing inhibition to the left excitatory pool leads to more right choices and vice versa. **(h,i)** Firing rates of PMd neurons for PREF direction reaches and NONPREF direction reaches aligned to checkerboard and movement onset. In Winner-take-all models, when one pool wins the firing rate of the other pool is suppressed due to lateral inhibition. In PMd, when the PREF direction wins the firing rate of the NONPREF direction decreases (blue shaded region in i).

**Figure S14:**
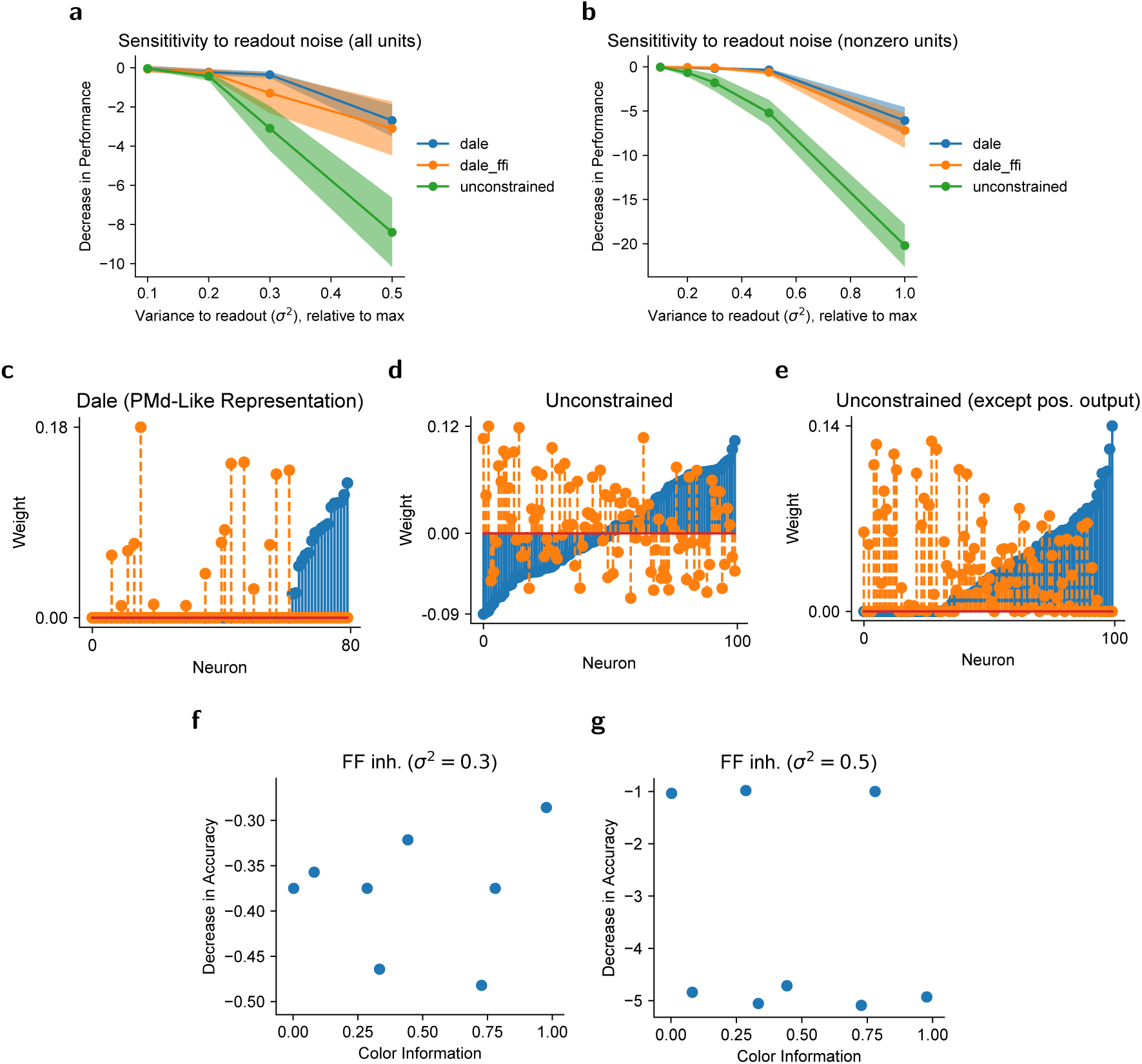
Potential multi-area computational advantage. **(Top Row)** Sensitivity to isotropic readout noise added to the output weights. **(a)** Noise added to all units in output (even the zero weights). **(b)** Noise only added to nonzero units. **(Middle Row)** Readout weights for left (dashed orange) and right (blue) reaches. **(c)** Readout weight with Dale’s Law enforced, (**d**) Readout weights in unconstrained networks. (**e**) Readout weights in unconstrained but ensuring positive outputs. **(Bottom Row)** No correlation between robustness to noise and usable color information across random initializations for networks with 10% feedforward inhibition, where after training some networks had color information (Fig. 5b). We used a noise perturbation to each unit of variance **(f)** *σ*^2^ = 0.3 and **(g)** *σ*^2^ = 0.5.

**Figure S15:**
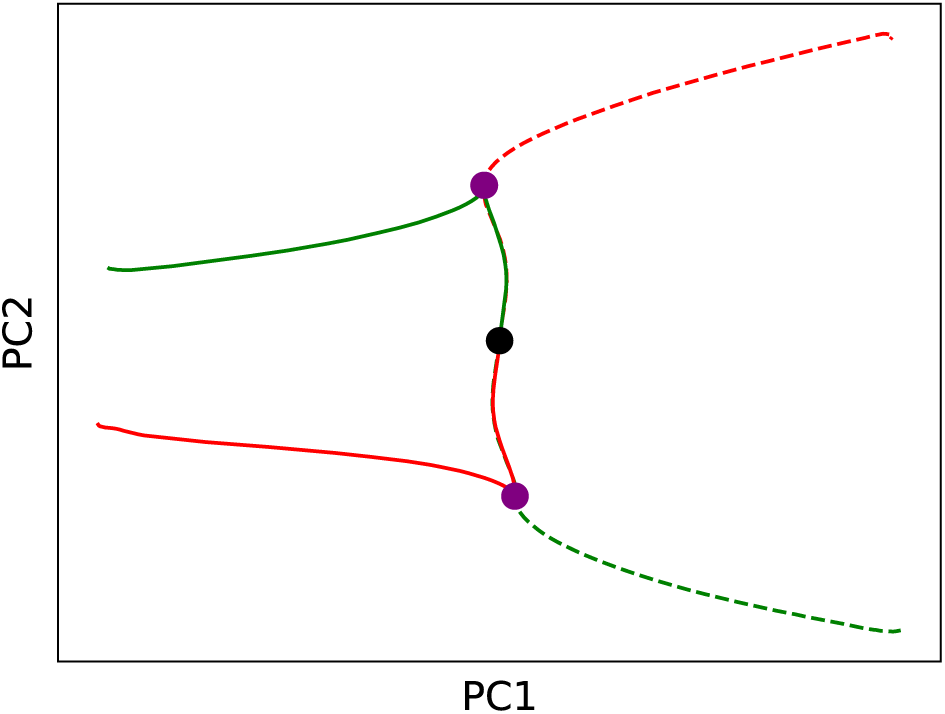
PCA of single area network (analogous to Fig 3d for a three-area network). Black circle denotes Checkerboard onset, and Purple circle denotes target onset. Variance numbers are 60.5%, 37.1% of PC1 and PC2 respectively. These PCs do not resemble those of PMd (Fig. 3f).

## Supplementary Note: Mutual Information Estimation

The entropy of a distribution is defined as

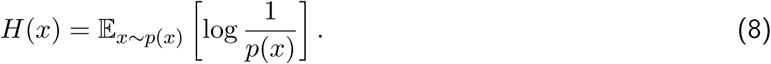

The mutual information, *I*(*X*; *Y*), can be written in terms on an entropy term and as conditional entropy term:

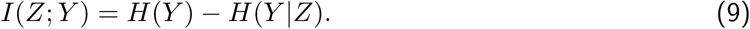

We want to show that the usable information lower bounds the mutual information:

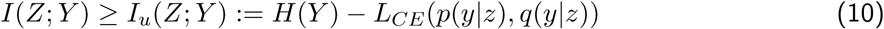

It suffices to show that:

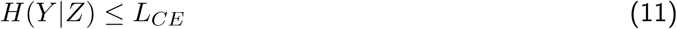

where *L_CE_* is the cross-entropy loss on the test set. For our study, *H*(*Y*) represented the known distribution of output classes, which in our case were equiprobable.

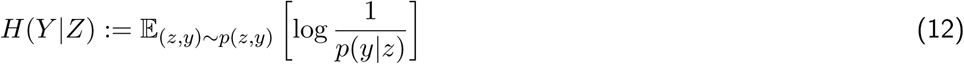

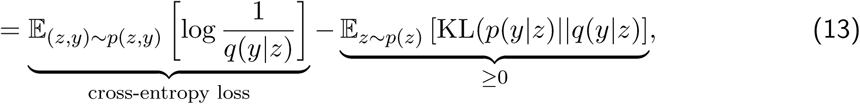

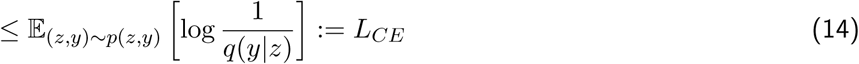

To approximate *H*(*Y* |*Z*), we first trained a neural network with cross-entropy loss to predict the output, *Y*, given the hidden activations, *Z*, learning a distribution *q*(*y*|*z*). The KL denotes the Kullback-Liebler divergence. We multiplied (and divided) by an arbitrary variational distribution, *q*(*y*|*z*), in the logarithm of equation 12, leading to equation 13. The first term in equation 13 is the cross-entropy loss commonly used for training neural networks. The second term is a KL divergence, and is therefore non-negative. In our approximator, the distribution, *q*(*y*|*x*), is parametrized by a neural network. When the distribution *q*(*y*|*z*) = *p*(*y*|*z*), our variational approximation of *H*(*Y* |*Z*), and hence approximation of *I*(*Z*; *Y*) is exact ^17,54,55^.

In the paper, we additionally report the accuracy of the neural network on the test set. This differs from the cross-entropy in that the cross-entropy incorporates a weighted measure of the accuracy based on how “certain” the network is, while the accuracy does not.

## Acknowledgments

MK was supported by the National Sciences and Engineering Research Council (NSERC). CC was supported by a NIH/NINDS R00 award R00NS092972, NIH/NINDS R01 NS122969, NIH/NINDS R01NS121409 the Moorman-Simon Interdisciplinary Career Development Professorship from Boston University, the Whitehall foundation, and the Young Investigator Award from the Brain and Behavior Research Foundation. JCK was supported by NSF CAREER 1943467, NIH DP2NS122037, the Hellman Foundation, and a UCLA Computational Medicine AWS grant. We gratefully acknowledge the support of NVIDIA Corporation with the donation of the Titan Xp GPU used for this research. We thank Laura Driscoll for helpful comments on the manuscript as well as Krishna V. Shenoy and William T. Newsome for helpful discussions on earlier versions of these results. We also thank Krishna V. Shenoy for kindly allowing us to use the PMd data collected by Dr. Chandrasekaran when he was a postdoc in the Shenoy Lab.

## Author contributions

MK, JCK and CC conceived of the study. MK and JCK trained RNNs and analyzed networks. MK performed the multi-area computation analyses. DX, EF, and NH assisted with various analyses. CC collected experimental data in PMd in Prof. Shenoy’s lab. TW, NC, and EKL trained animals and collected DLPFC data in the lab of CC. TW performed various analyses on the neural data. YL contributed by curating data and code for further analysis. MK, CC and JCK wrote the manuscript.

